# Data-poor stock assessment of fish stocks co-exploited by commercial and recreational fisheries: applications to pike (*Esox lucius*) in the western Baltic Sea

**DOI:** 10.1101/2021.01.20.427466

**Authors:** Rob van Gemert, Dieter Koemle, Helmut Winkler, Robert Arlinghaus

**Affiliations:** Department of Biology and Ecology of Fishes, Leibniz Institute of Freshwater Ecology and Inland Fisheries, Müggelseedamm 310, 12587 Berlin, Germany; Department of Aquatic Resources, Institute of Freshwater Research, SLU-Aqua, Drottningholm, Sweden; Institute of Biodiversity and Zoology, University of Rostock, Rostock, Germany; Division of Integrative Fisheries Management, Faculty of Life Sciences, Humboldt-Universität zu Berlin, Philippstraße 13, Haus 7, 10115 Berlin, Germany

**Keywords:** Catch-only models, superensemble models, mixed-use fisheries, stock status, freshwater stocks, coastal stocks

## Abstract

Information on catch and effort of recreational angling in mixed-use fisheries (co-exploited by commercial and recreational fishers) is often scarce, preventing the application of data-rich stock assessments typically performed for industrialized commercial fisheries. Here, we show how data-poor stock assessment methods developed for marine fisheries, particularly a class of models labelled as “catch-only” models (COMs), offer a possible solution. As a case study, we use COMs to assess a northern pike stock around the German Baltic island of Rügen. We fit multiple COMs to a time-series of total pike removals, and use their outputs as explanatory variables in superensemble models. We conclude that the stock is fully exploited and currently declining. Our study highlights the potential for using COMs to determine status of previously-unassessed coastal and freshwater stocks facing recreational fishing pressure, and demonstrates how incorporating recreational removals is crucial for achieving reliable insights into the status of mixed-use stocks.

## Introduction

The management of mixed-use fisheries, i.e., fisheries that are co-exploited by commercial and recreational fishers, poses many challenges. For instance, commercial and recreational fisheries will often have different management objectives (Arlinghaus et al., 2019; Ahrens et al., 2020), and the disparity in aspirations and behaviours increases with the diversity of stakeholders (Mardle et al., 2004; Pascoe et al., 2009). The sustainable management of mixed-use fisheries requires addressing both its commercial and recreational components, because the combined action of both sectors is responsible for the total fishing mortality induced on a stock (Berkes, 1985; Post et al., 2002; Cooke & Cowx, 2004).

A common precondition for sustainability in fisheries is the existence of regular stock assessments (Melnychuk et al., 2017). Stock assessments employ a variety of methods depending on the data available. The most reliable “data-rich” methods (such as virtual population analysis and statistical catch-at-age models) require data on catch, effort, and the age/length/weight composition, preferably from both the fishery users as well as from independent scientific surveys (Hilborn & Walters, 1992; Hart & Reynolds, 2002). The management of industrial commercial fisheries appears to become increasingly effective due to the presence of frequent and high quality stock assessments, with many assessed stocks showing rebuilding from previously overfished states (Hilborn et al., 2020). However, gathering the required data for such assessments is costly (Mangin et al., 2018), and therefore high quality data to pursue stock assessments are rarely available for many small-scale commercial (Andrew et al., 2007; Graaf et al., 2015; Prince & Hordyk, 2019) and recreational fisheries (Post et al., 2002; Arlinghaus et al., 2019). As a result, mixed-use fisheries can be expected to often face a severe lack of data, preventing the application of standard stock assessment practices common to industrial commercial fisheries.

When there is insufficient data available for performing a traditional data-rich stock assessment, the fishery is usually referred to as data-poor or data-limited (Prince & Hordyk, 2019). Many small-scale commercial and recreational fisheries are characterized as such, with aggregated catch or landings data often being the only form of data available (Vasconcellos & Cochrane, 2005; Newman et al., 2015). An alternative to a stock assessment is to infer stock status from trends in catch data, as is done in stock-status plots (Grainger & Garcia, 1996; Froese & Kesner-Reyes, 2002; Pauly, 2007). However, because catches do not necessarily track changes in underlying biomass, such catch-based methods can result in incorrect conclusions (Branch et al., 2011; Daan et al., 2011; Carruthers et al., 2012). To overcome this problem and still be able to make predictions on stock status using aggregated catch data, more sophisticated models have been developed that either rely on underlying population dynamics models (Vasconcellos & Cochrane, 2005; Martell & Froese, 2013), or statistical correlations with data-rich assessed stocks (Costello et al., 2012; Zhou et al., 2017). Here, we call these models catch-only models (COMs).

COMS designed to estimate stock status time-series can be divided into two broad categories: mechanistic and empirical COMs (Free et al., 2020). Mechanistic COMs fit a population dynamics model to the catch data and make assumptions regarding parameter values to make up for the lack of other data. Mechanistic COMs include models such as CMSY (catch maximum sustainable yield; Froese et al., 2017) and SSCOM (Thorson et al., 2013). Empirical COMs use information from data-rich assessed stocks to find statistical associations between catch, stock status, and other covariates. Empirical COMs include models such as mPRM (Rosenberg et al., 2014) and zBRT (Zhou et al., 2017). COMs are not as accurate in predicting stock status as data-rich statistical catch-at-age models, but they offer a promising alternative when the absence of some data prevents a full data-rich assessment (Free et al., 2020).

In addition to using quantitative models to determine the status of data-poor stocks, assessments based on traditional ecological knowledge of local residents have been performed as well (Berkes et al., 2000). Individual perceptions can conflict with scientific findings (O’Donnell et al., 2010; Daw et al., 2011), but for many data-poor stocks such local knowledge can be one of the few sources of information on the development of the stock (Johannes, 1998). Furthermore, studies that have compared traditional ecological knowledge with independent stock assessments have often found that model outcomes align with local understanding (Neis et al., 1999; Aswani & Hamilton, 2004) and that local users can approximate scientific understanding of ecological relationships in fish stocks (Aminpour et al., 2020). Thus, the knowledge of local residents can be used to see whether scientific and stakeholder perspectives agree.

In this study, we explore the usage of COMs to assess the status of a data-poor mixed-use single-species fishery, using a coastal lagoon pike (*Esox lucius*) stock in the western Baltic Sea in Germany as a case study. The pike stock around the island of Rügen is targeted by both recreational and commercial fishers, but regular stock assessments are lacking and disparate perspectives about stock status have emerged among stakeholders that contribute to local conflict (Vogt, 2020). To help solve these issues, we assessed the status of this coastal pike stock using seven different COMs, and used a superensemble model approach to account for individual model biases. We show the importance of using multiple different COMs for assessing stock status, and reveal how including catch data from both its recreational and commercial components is crucial for the assessment of a mixed-use fishery. Furthermore, we show that surveys among local stakeholders can be used to increase the confidence in model results. Using a practically-relevant example of an ongoing management dilemma in a mixed-use coastal fishery, we demonstrate how COMs can be used as an initial method for the assessment of data-poor mixed-use stocks, being aware that it is not a perfect substitute for more data-rich approaches.

## Materials & Methods

### Study area and data

We studied the pike stock around the German island of Rügen, which is located in the western Baltic Sea (Figure 1). There are multiple lagoon-type brackish water bodies located around this island, which are connected to the Baltic Sea (Schubert & Telesh, 2017). These lake-like water bodies vary in salinity from nearly fresh to nearly that of the neighboring Baltic Sea (Placke et al., 2018) and exhibit salinities below 14 PSU year-round, i.e., mesohaline brackish conditions (Schumann et al., 2006; Schiewer, 2008). Although pike is a freshwater fish, the species is able to tolerate the brackish water present in the lagoon waters and is known to successfully spawn and recruit in the brackish conditions around Rügen (Möller et al., 2019).

**Figure 1:**
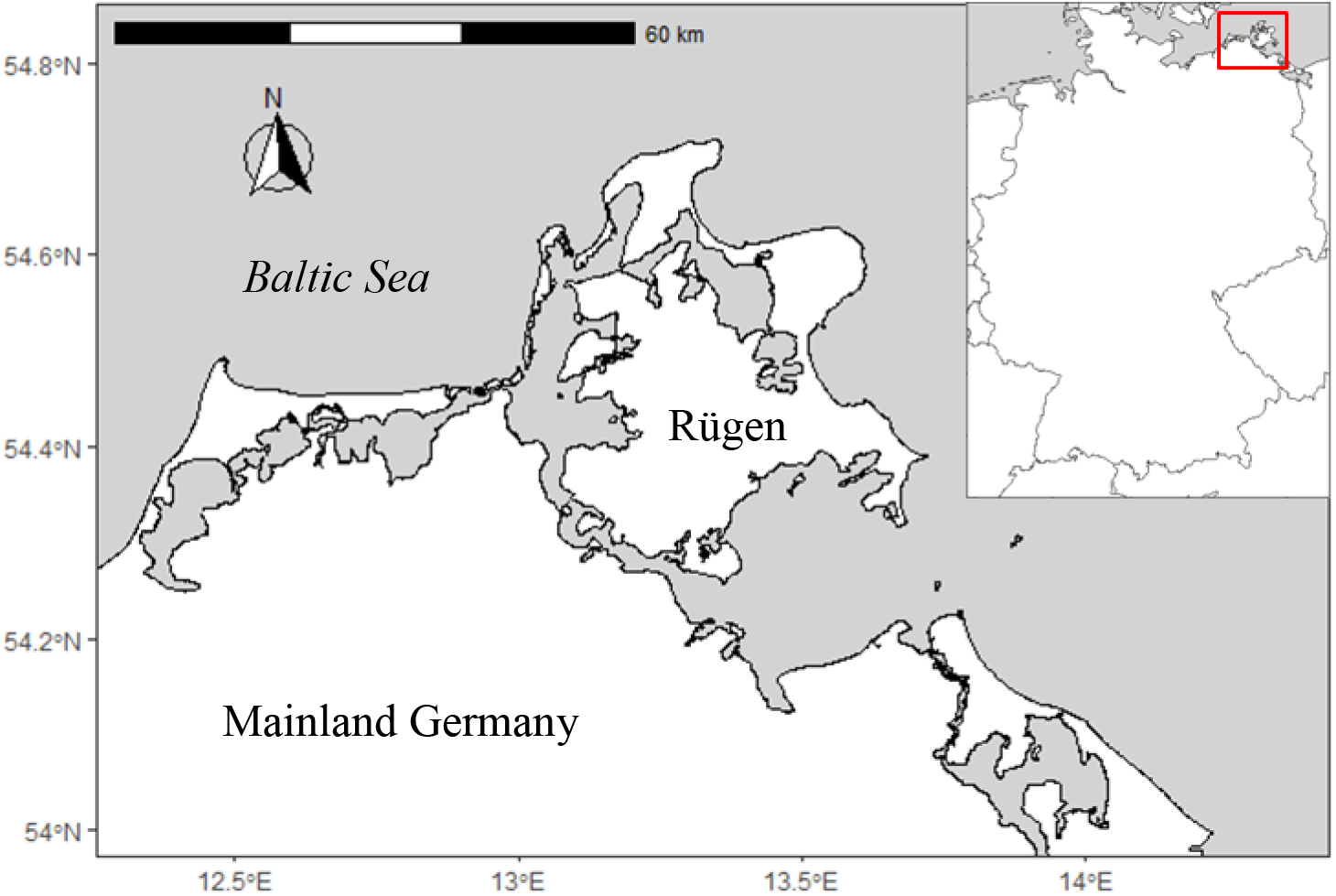
A map of the German Baltic island of Rügen, also indicating its location within Germany.

The pike stock around the lagoons of Rügen has been exploited by small-scale coastal commercial fishers since at least the late 19th century (Winkler, 1989; Winkler & Debus, 2006), and likely long before. For much of the 20th century, the pike was a target for commercial fishers, helped by the guaranteed price per unit weight that was maintained by the East Germany (GDR) government (Döring et al., 2020). However, after the German reunification, the pike has become a bycatch species for most coastal commercial fishing enterprises and is today the primary target for only a small number of dedicated commercial pike fishers. However, the species continues to be captured in gill nets, fyke nets, and with long-lines up to the present day.

Recreational angling was practiced in the Rügen lagoons during GDR times, and has been increasing in popularity after the German reunification (Figure A5, Appendix A). Pike is a particularly popular fish for recreational fishers around Rügen, especially for non-resident anglers, likely due to its relatively large size in the lagoon ecosystems, but is also a valued target of resident anglers (Weltersbach et al., in press).

### Catch time-series data

A time-series of annual commercial pike landings from 1976-2018 from the lagoons around Rügen was obtained from the state office for agriculture, food safety, and fisheries (LALLF) of the German state Mecklenburg-Vorpommern (M-V). Furthermore, commercial removals from 1969-1975 and from 1955-1968 were extracted from Winkler (1991) and from records generated from annual official fisheries reports published by the Institut für Hochseefischerei und Fischverarbeitung Rostock of the former GDR, respectively.

A time-series of recreational removals was not available for the Rügen pike stock, as recreational removals are not actively monitored in the area by public authorities. However, given the popular recreational fishery for Rügen pike, we considered it important to include recreational removals in the time-series of total removals. We reconstructed recreational removals according to the guidelines provided by Zeller *et al*. (2006). We used data from two telephone-diary studies among recreational fishers performed in the region (Dorow & Arlinghaus, 2011; Lucas, 2018; Weltersbach et al., in press) as anchor points. These studies estimated total pike removals in the Rügen area from a random sample of participating recreational fishers for the years 2006/2007 and 2014/2015. From these anchor points, and using additional quantitative data such as proxies for angling effort, we inter-and extrapolated recreational removals for the rest of the time-series.

The reconstruction of recreational removals of the Rügen pike (detailed in Appendix A) can be summarized as follows. For the year 2007, data were available on resident angler harvest and catch-and-release rate, and the number of angling trips taken in the Rügen area by resident anglers (Dorow & Arlinghaus, 2011). Comparable data were available for 2015, not only for resident but also for tourist anglers (Lucas, 2018; Weltersbach et al., in press). These anchor points were supplemented with time-series data on the annual number of resident fishing licenses issued in M-V as given by LALLF, the annual number of coastal recreational angling licenses issued in M-V as given by LALLF, and the membership numbers of the East German angling association (DAV) prior to the German reunification (VDFF, 1998). We then used the 2007 and 2015 data on resident angling trips together with the time-series on coastal recreational angling licenses to estimate a 1991-2018 time-series on resident angling trips, assuming a linear increase in trips per license for resident anglers between 2007 and 2015. Similarly, we used the 2015 data on tourist angler trips together with the time-series on coastal recreational angling licenses to estimate a 1991-2018 time-series on tourist angling trips, assuming a constant number of trips per license for tourist anglers based on the 2015 data. Then, we used DAV membership data to extrapolate this recreational effort time-series back to 1955. Next, using available time-series data on commercial Rügen pike removals and the annual number of commercial fishing vessels registered in the area as given by the European Fleet Register, we estimated a commercial catch-per-unit-effort (CPUE) time-series (landings per boat), assuming a constant commercial effort in the years for which we lacked data (1955-1991). Then, we used this commercial CPUE time-series together with the 2007 and 2015 data on recreational fisher catch rates to estimate a 1991-2018 time-series of recreational catches, assuming constant proportionality between recreational and commercial CPUE for all years. We subsequently estimated resident and tourist removals by accounting for the release rates of 2007 and 2015, assuming a linear decrease in resident release rate between 2007 and 2015 and assuming that tourist release rate remained constant to its 2015 value, and furthermore assuming a release mortality for pike of 7.8% (Hühn & Arlinghaus, 2011). Lastly, we used the reconstructed recreational removals of 1991 to extrapolate recreational removals back to 1955, by assuming a constant proportionality between recreational and commercial CPUE.

### Models

First, we used a suite of different COMs to estimate current status of the Rügen pike stock in an ensemble model approach. We then inserted the results of these models into several different ‘trained’ statistical models, following the superensemble model approach as described by Anderson *et al*. (2017), providing an estimate of current stock status. Lastly, we used the outcome of the superensemble analysis to assign weights to COM time-series estimates of biomass and fishing mortality, providing an estimate of past stock status. The analysis was performed in R version 3.6.1 (R Core Team, 2019), using the `datalimited` (Anderson et al., 2016) and `datalimited2` (Free, 2018) packages for the COMs, and the `randomForest` (Liaw & Wiener, 2002) and `gbm` (Greenwell et al., 2019) packages for the superensemble model.

### Catch-only models (COMs)

We fitted seven individual COMs to the reconstructed removal data of the Rügen pike stock. We used the COMs that had their performance tested by Free et al. (2020). These included five COMs that fit a population dynamics model, and two COMs that find statistical associations using data-rich assessed stocks. Each of the COMs returned an estimate of *B/B_MSY_* over the course of the catch time-series, including a 95% confidence interval. Furthermore, parameters and reference points returned by some, but not all, COMs include fishing mortality *F*, fishing mortality that gives MSY *F*_MSY_, biomass *B, B*_MSY_, MSY, intrinsic population growth rate *r*, and population carrying capacity *k*. The COMs that were used were Catch-MSY (Martell & Froese, 2013), CMSY (Froese et al., 2017), COM-SIR (Vasconcellos & Cochrane, 2005), SSCOM (Thorson et al., 2013), mPRM (Rosenberg et al., 2014), OCOM (Zhou et al., 2018), and zBRT (Zhou et al., 2017). A brief explanation of each model is provided in Appendix B.

### Superensemble models

The different COMs that we fitted to our data yield different results, owing to their inherent biases due to different methods and assumptions (Free et al., 2020). To try and resolve these potential discrepancies, we combined the results of the different COMs in a superensemble model approach. For this, we used the approach described by Anderson et al (2017). Firstly, for each COM, we took the estimated values for mean and slope of *B/B*_MSY_ and computed an average for the last 5 years of data. Second, we inserted these COM averages as covariates into three different statistical models (here called superensemble models) that were trained on stocks with known status, thereby obtaining a superensemble estimate of the mean and slope of *B/B*_MSY_ for the last 5 years. The three statistical models that we used for this were a linear model, a random forest, and a boosted regression tree.

To train the three superensemble models with known data, we used simulated exploitation time-series for 5,760 different hypothetical stocks. For this, we used the exploitation time-series simulated by Rosenberg et al. (2014), which varied in both life-history parameters and fishing regime, and contained both process and observation error. Then, for each hypothetical stock, we fitted each of the seven COMs to its simulated catch time-series, giving seven estimates of a time-series for *B/B*_MSY_ for each stock. Next, for each of these time-series, we took the mean and slope of *B/B*_MSY_ and calculated an average over the last 5 years of data. Lastly, we trained the statistical models, using either the mean or the slope of recent *B/B*_MSY_ estimated by the seven COMs for all 5760 simulated stocks as seven independent variables, and the associated true mean or slope of recent *B/B*_MSY_ of all simulated stocks as the dependent variable. To prevent overfitting of the random forest and boosted regression tree models, we used the caret package (Kuhn, 2020) in R to identify the optimal parameter combination for both the *B/B*_MSY_ mean and slope model fits. Using a 10-fold cross validation repeated 10 times, optimal parameter combinations were identified as those that resulted in the smallest root-mean-square deviation. Following the results of this tuning procedure (Appendix B), we fitted both random forest models for mean *B/B*_MSY_ and its slope using 3 randomly selected predictors and 1000 trees, we fitted the boosted regression tree model for mean *B/B*_MSY_ using 9500 trees, an interaction depth of 10, and a shrinkage of 0.005, and we fitted the boosted regression tree for *B/B*_MSY_ slope using 7000 trees, an interaction depth of 10, and a shrinkage of 0.005, keeping all other parameters to their default setting.

After the superensemble models have been trained with known data, they were used to estimate the status of the Rügen pike stock. For this we used the outcomes of the seven COMs (estimates of the mean value and mean slope of *B/B*_MSY_ of the Rügen pike stock over the last 5 years of data) as the independent variables, whilst retaining the values of the regression coefficients estimated in the training of the superensemble models. In this way, each superensemble model provided an estimate of the mean and slope of *B/B*_MSY_ for the Rügen pike over the last 5 years of data. To study the importance of incorporating recreational removals into the total removals time-series, we repeated this process using only the commercial landings of pike as input to the COMs and the subsequent superensembles.

### Estimating F and F_MSY_

Aside from a *B/B*_MSY_ time-series, four mechanistic COMs (Catch-MSY, CMSY, COM-SIR, and OCOM) also estimate *F* and *F*_MSY_. We used these COMs’ weighted means of *B, B*_MSY_, *F*, and *F*_MSY_ to construct a Kobe plot, showing the recent trend in stock status relative to *F*_MSY_ and *F*_MSY_. To construct a weighted mean of each of these variables, we assigned COM-specific weights. These weights were based on each COM’s percentage error of its estimate of mean *B / B*_MSY_ over the last 5 years, compared to the mean of the superensemble estimates. Percentage error *p*_error_ of COM *i* was calculated as:

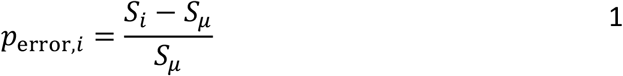

where *S* is the COM estimate of mean *B/B*_MSY_ over the last 5 years, and *S_μ_* is the mean of the superensemble estimates of mean *B/B*_MSY_ over the last 5 years. Next, weight *w* of COM *i* was calculated as the reciprocal of the absolute value of *p*_error_:

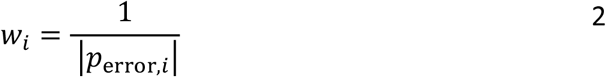

Thus, if a COM estimate of *B/B*_MSY_ had a larger deviance from the superensemble model average, it received a smaller weight. Weights were subsequently normalized according to:

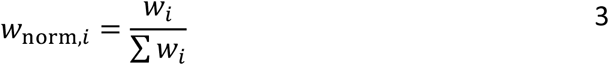

Normalized weight *w*_norm_ of a COM was then used to calculate weighted means of *B, B*_MSY_, *F*, and *F*_MSY_ for each of the COMs that estimated these values.

### Sensitivity tests

Sensitivity tests were performed to analyze the sensitivity of the models to alternative parameter values, and to test a number of the assumptions made in our reconstruction of recreational removals. The methodology and results of these tests are described in detail in Appendix D. In summary, we first performed an elasticity analysis to test the sensitivity of the model results to changes in COM parameter values. We varied each parameter by 50%, and considered a model sensitive to a given parameter when the model result deviance from its base run estimate was greater than 50%. Second, we looked at the sensitivity of the model results to assumptions made during the reconstruction of the recreational removals. We reconstructed the recreational removals time-series through various alternative methods, ran the COMs and superensemble models with the resulting time-series of total removals, and examined the deviance of each model’s result from its base run.

### Stakeholder perceptions of stock trends

To gain insight in how different stakeholders perceived the development of the Rügen pike stock, we constructed a short questionnaire and distributed it among the key stakeholder groups (anglers, angling guides, commercial fishers, non-governmental organizations, and fisheries agency staff). Among other things, we specifically asked respondents how they perceived the stock of pike to have changed within the time-frame between today and the first time they fished at the Rügen lagoons. The same question was asked regarding the stock development of pike greater than one meter in length. Responses were measured on a five-point Likert scale from “strong decrease” to “strong increase”.

The survey was administered through a snowball technique to both resident and non-resident anglers as well as angling guides, fishers, and other stakeholders. Data were further collected on an angling exhibition in Rostock in June 2019, and through local angling guides. The total sample size numbered 258 observations. The resulting data are not representative for the population-level perceptions, but allowed us to gather initial insights for the most heavily engaged stakeholders, and to compare stock trends derived from the COMs with stakeholder perspectives.

## Results

The seven COMs predicted different historic patterns in the *B/B*_MSY_ trend of the pike stock in the Rügen area (Figure **4**). Notably, zBRT predicted a severely depleted stock status in the 1970s and 1980s, while most other COMs predicted a status of *B/B*_MSY_ remaining around or above 1 for the majority of the time-series. Similarly, estimates of current *B/B*_MSY_ varied, with two COMs estimating a current *B/B*_MSY_ greater than 1, one COM estimating it lower than 1, and four COMs estimating it to be around 1. However, all COMs consistently estimated a decline in *B/B*_MSY_ in recent years.

**Figure 2:**
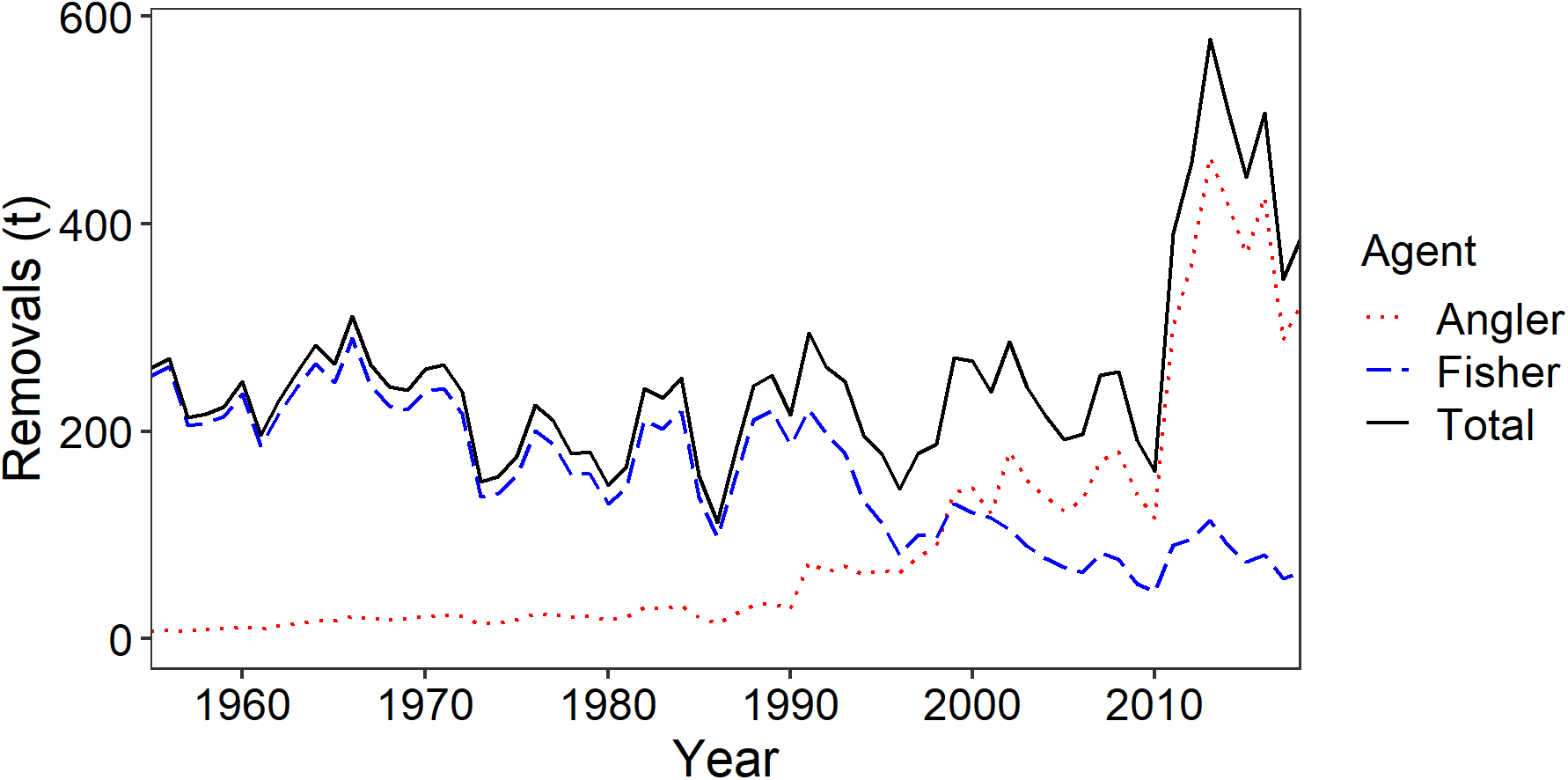
Total removals by both commercial and recreational fishers in the Rügen area over time (solid black). Shown too are removals by commercial fisheries over time (dashed blue) and reconstructed removals by recreational fishers over time (dotted red).

**Figure 3:**
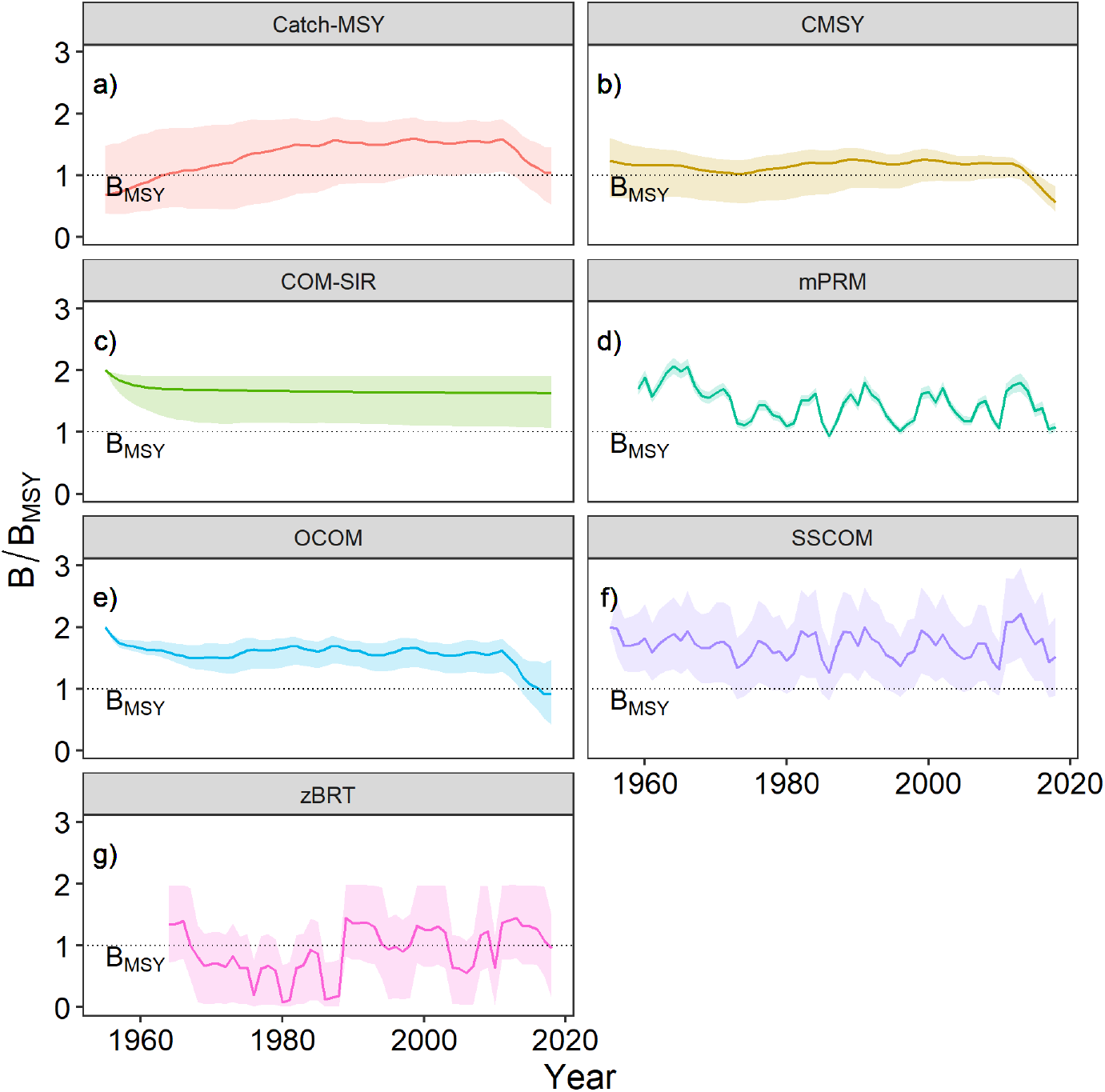
*B/B*_MSY_ trends as estimated by seven individual COMs: Catch-MSY (a), CMSY (b), COM-SIR (c), mPRM (d), OCOM (e), SSCOM (f), and zBRT (g). Shaded areas indicate 95% confidence intervals, the dotted line indicates *B*_MSY_.

**Figure 4:**
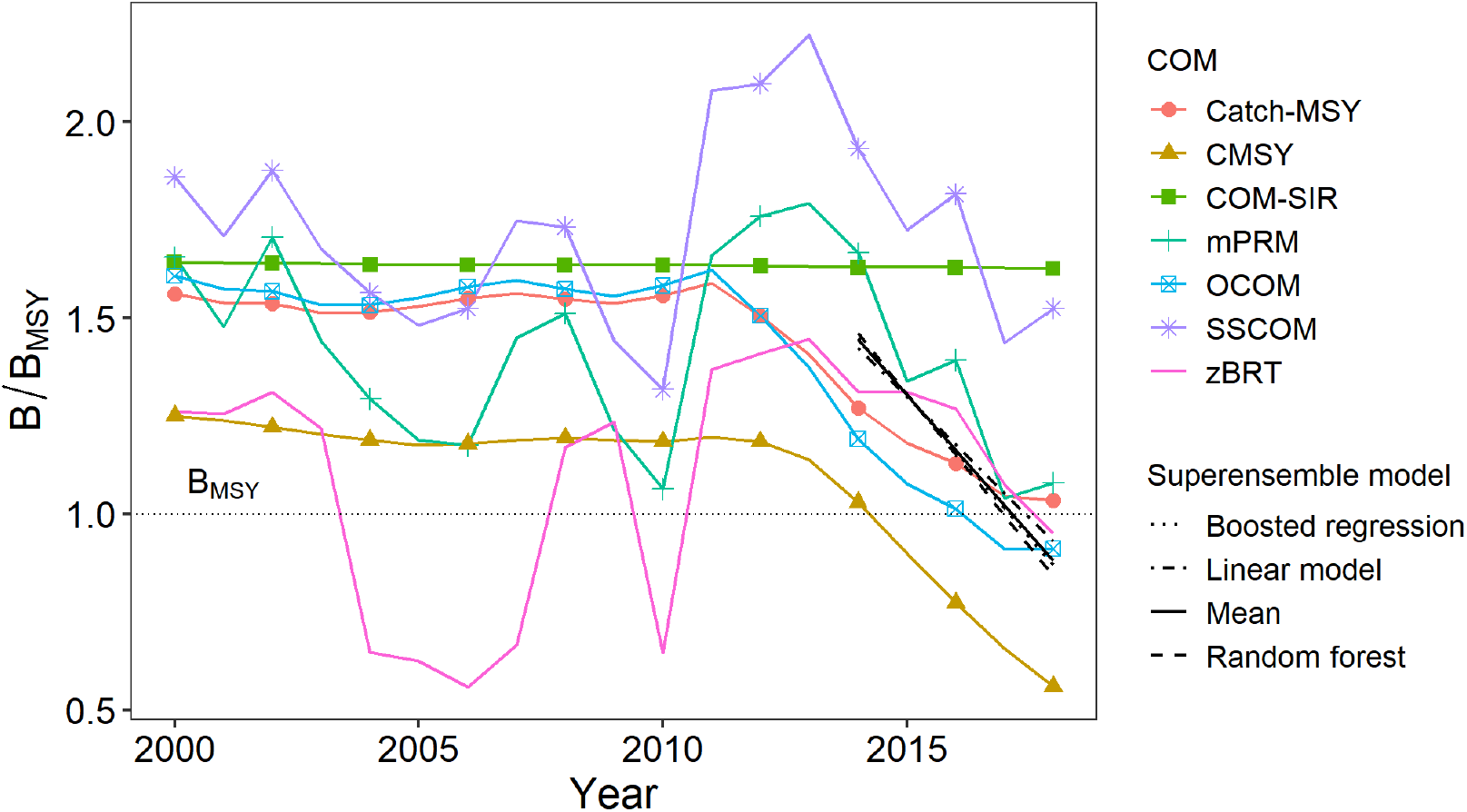
Results of the superensemble model estimates of mean and slope of *B/B*_MSY_ over the last 5 years of data, including their overall mean, overlaid on a truncated time-series of individual COM results. The horizontal dotted line indicates *B*_MSY_.

The different superensemble models predicted similar values for mean *B/B*_MSY_ in the past 5 years, and all predicted a negative slope of recent *B/B*_MSY_ (Table 1). Although all superensemble models predicted a 5-year mean of recent *B/B*_MSY_ above 1, extrapolating this mean with each superensemble’s estimated slope *B/B*_MSY_ suggested that current *B/B*_MSY_ of the Rügen pike stock is around or even slightly below 1 (Figure **5**). Thus, based on the superensemble model results, the Rügen pike stock is fully exploited, but may also be slightly growth overfished when judged by MSY.

**Figure 5:**
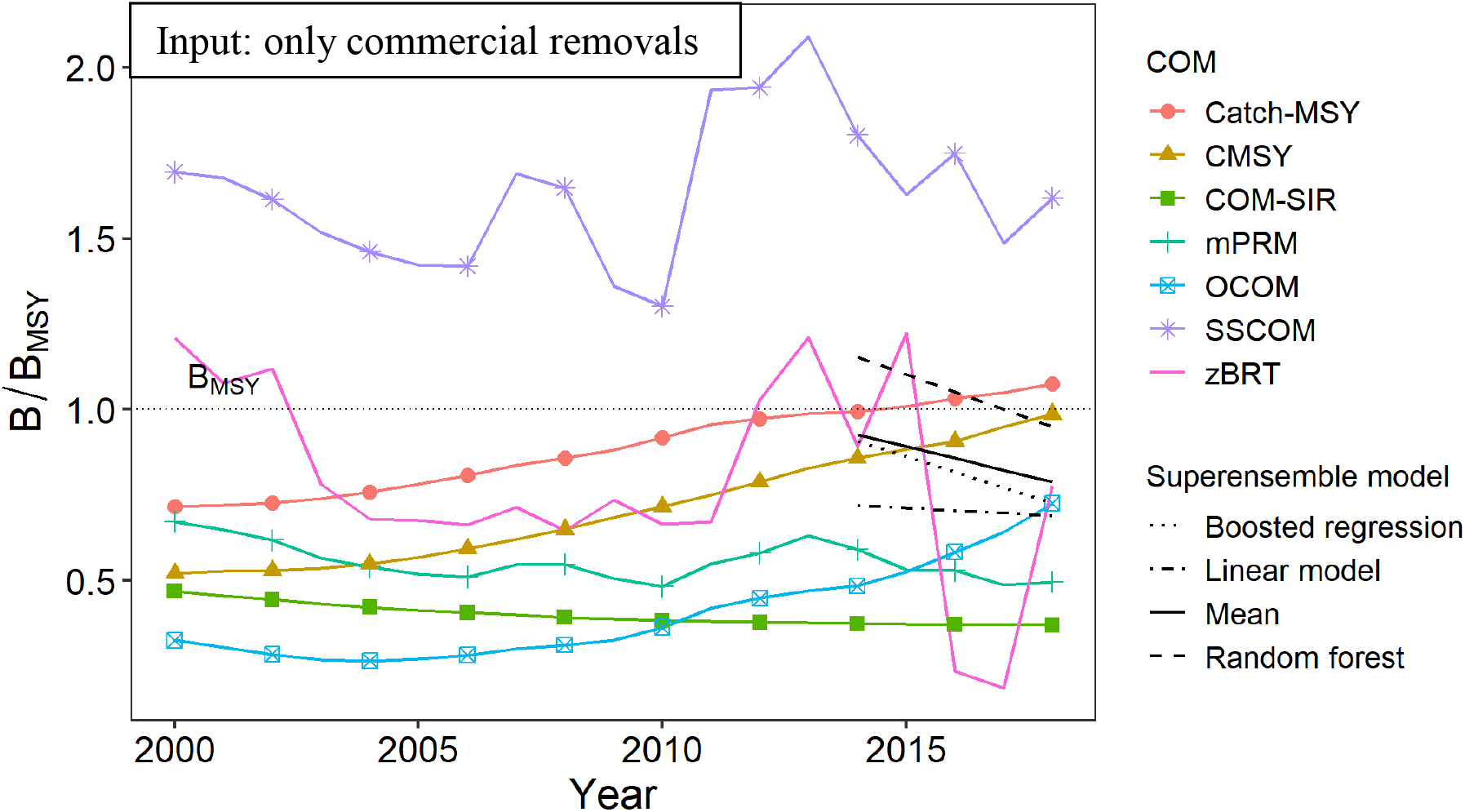
Model output when only the commercial removals time-series is used as input, and the recreational removals time-series is left out. Shown are the subsequent results of the superensemble model estimates of mean and slope of *B/B*_MSY_ over the last 5 years of data, including their overall mean, overlaid on a truncated time-series of individual COM results. The horizontal dotted line indicates *B*_MSY_.

**Table 1:** COM and superensemble model estimates of the mean value of *B/B*_MSY_ in the last 5 years of data, and the average slope of *B/B*_MSY_ in the last 5 years of data. Shown in bold are the averages of all COMs and all superensemble models, respectively.

| Model | Mean | Slope |
| --- | --- | --- |
| *COM* |  |  |
| Catch-MSY | 1.13 | -0.0622 |
| CMSY | 0.784 | -0.121 |
| COM-SIR | 1.63 | -0.000620 |
| mPRM | 1.30 | -0.147 |
| OCOM | 1.02 | -0.0762 |
| SSCOM | 1.69 | -0.105 |
| zBRT | 1.18 | -0.100 |
| **COM average** | **1.21** | **-0.0875** |
| *Superensemble* |  |  |
| Boosted regression | 1.16 | -0.144 |
| Linear model | 1.18 | -0.123 |
| Random forest | 1.15 | -0.155 |
| **Superensemble average** | **1.16** | **-0.141** |

When recreational removal data was left out of the analysis, and the models were provided with only the commercial catch data, clear differences from the original estimates of the catch-only and superensemble models could be observed (Figure **6**). When not considering recreational fishers, the pike stock appeared in a much poorer and highly overfished state than when recreational removals were included.

**Figure 6:**
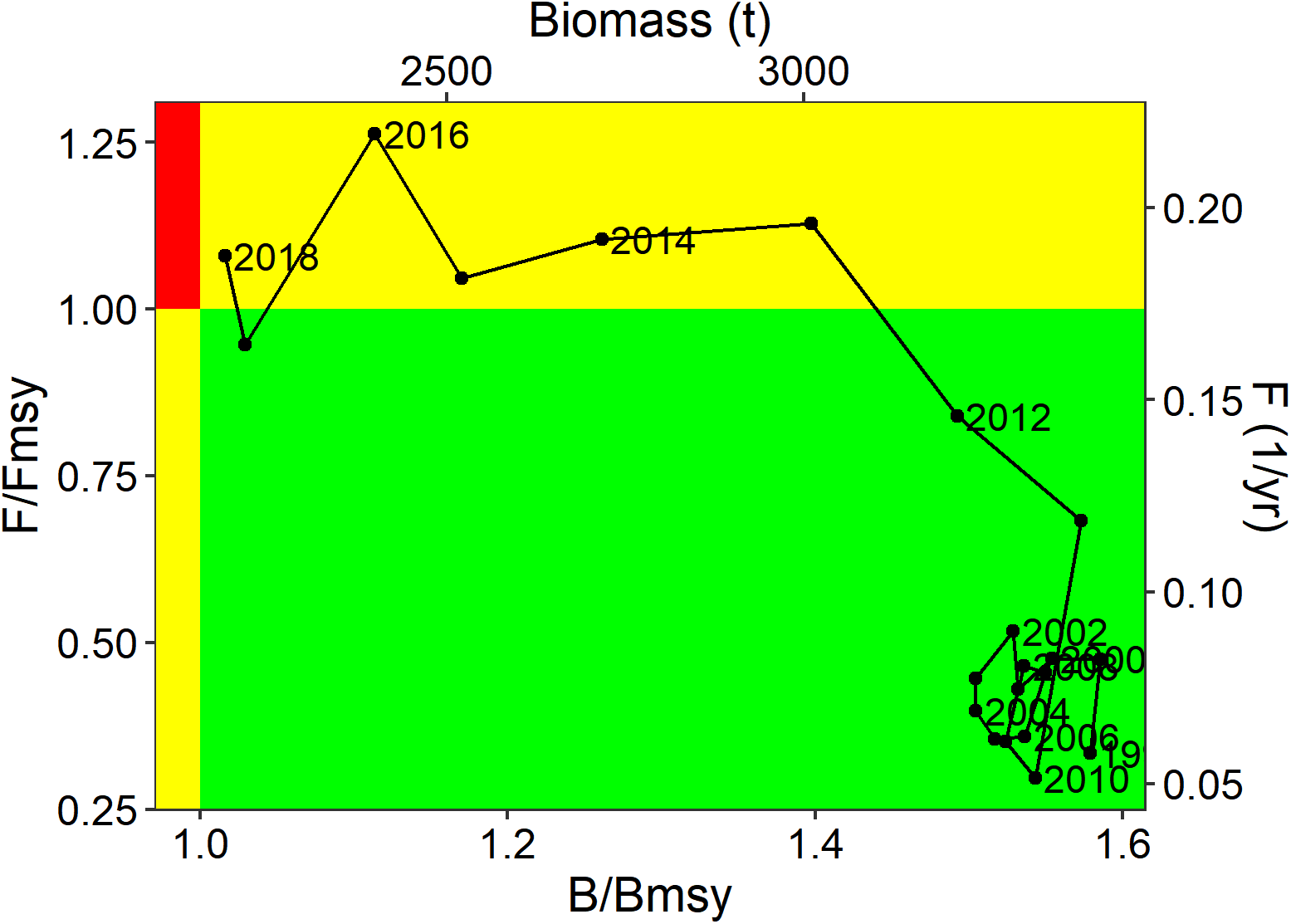
Kobe plot showing the weighted averages of *B,B/B*_MSY_, *F*, and *F/F*_MSY_ for the years 1998 to 2018. The green area indicates healthy stock status, the yellow areas indicate that the stock is either overfished (*B/B*_MSY_ < 1) or subject to overfishing (*F/F*_MSY_ > 1), and the red area indicates that the stock is both overfished and subject to overfishing.

The model weights for the calculations of weighted mean values of *B, B/B*_MSY_, *F*, and *F/F*_MSY_ are given in Table 2. The Catch-MSY model was by far assigned the greatest weight. The resulting weighted averages of *B, B/B*_MSY_, *F*, and *F/F*_MSY_ were visualized as a Kobe plot (Figure **7**), and indicate that the stock had a healthy status and was not experiencing overfishing up until 2012, after which overfishing gradually reduced stock biomass to nearly below *B*_MSY_. Thus, the pike stock is currently experiencing overfishing, and is fully exploited. This result contrasts with the results of the superensemble models, which predict a current *B* state below *B*_MSY_. This discrepancy can be attributed to the large weight assigned to the Catch-MSY model, which estimates a current B state above *B*_MSY_ (Figure **4**). Using the model weights, estimates of current state of the Rügen pike stock, fisheries reference points, and life-history parameters were calculated from their individual COM estimates as summarized in Table 3.

**Figure 7:**
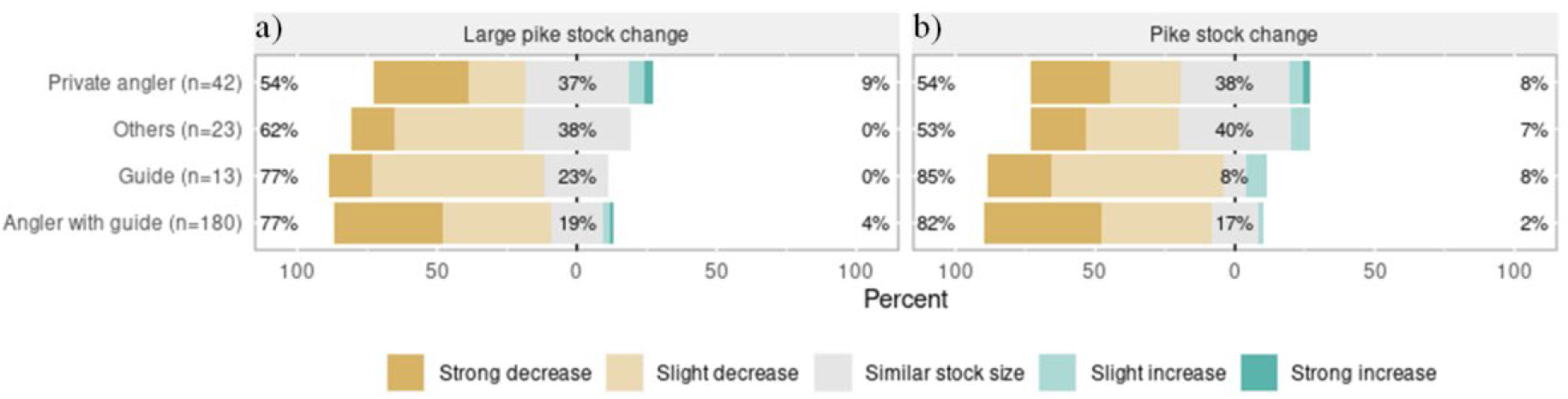
Stakeholder perceptions regarding the recent trends in the number of large pike of the Rügen stock (a) and the total size of the Rügen pike stock (b).

**Table 2:** Weights assigned to the listed COMs, used to calculate time-series of the weighted means of *B,B/B*_MSY_, *F*, and *F/F*_MSY_, as well as estimates of current state of the Rügen pike stock, fisheries reference points, and life-history parameters.

| Model | Normalized weight |
| --- | --- |
| Catch-MSY | 0.737 |
| CMSY | 0.0587 |
| COM-SIR | 0.0198 |
| OCOM | 0.0783 |

**Table 3:** Estimates of the 2018 state of the Rügen pike stock, fisheries reference points, and life-history parameters, including 95% confidence limits. Estimates are calculated as a weighted mean of four COMs: Catch-MSY, CMSY, COM-SIR, and OCOM.

| Estimate | Value | Lower limit | Upper limit |
| --- | --- | --- | --- |
| 2018 $F$ ($\text{yr}^{-1}$) | 0.18 | 0.084 | 0.52 |
| 2018 $B$ (t) | 2217 | 592 | 6914 |
| MSY (t) | 356 | 205 | 597 |
| $F_{\text{MSY}}$ ($\text{yr}^{-1}$) | 0.17 | 0.055 | 0.35 |
| $B_{\text{MSY}}$ (t) | 2154 | 1103 | 4734 |
| $r$ ($\text{yr}^{-1}$) | 0.347 | 0.109 | 0.699 |
| $k$ (t) | 4308 | 2207 | 9468 |

The various sensitivity analyses showed that the mean of the superensemble model estimates was generally robust to parameter settings and model assumptions (Appendix D). The elasticity test showed that, in general, the models were insensitive to changes in parameter values, with the greatest sensitivity being to changes in the prior of final year biomass range. Next, regarding the assumption of constant commercial effort before 1991, the results remained largely unchanged when different assumptions were made regarding the trend of commercial effort before 1991. Lastly, reconstructing recreational removals in a variety of different ways changed the mean of superensemble estimates of mean *B/B*_MSY_ over the last 5 years to only a limited degree, with a positive deviance of 20.8% being the highest among all different reconstructions, followed by a positive deviance of 14.2%.

From the stakeholder survey, when asked about their perceptions of the development of the Rügen pike stock, respondents mostly indicated a perceived decline over time in large pike of the Rügen stock as well as a perceived decline for the Rügen pike stock in general (Figure 8). This perceived decline was greatest among guides and those anglers who were out with a guide when interviewed. Thus, stakeholders agreed with the model results in estimating a recent decline of the pike stock.

## Discussion

Mixed commercial-recreational fisheries can be challenging to manage because commercial and recreational fisheries typically have divergent management objectives (Arlinghaus et al., 2019; Ahrens et al., 2020). In this context, lack of data can hinder the performance of standard stock assessments, which are an important instrument in the effective and sustainable management of fish stocks (Melnychuk et al., 2017; Hilborn et al., 2020). The application of COMs and superensemble models can be used to deliver insights into stock status when the only fisheries data available are observations of aggregated landings. Here, seven different COMs and three different superensemble models were applied in a case study to assess the status of pike in a German lagoon system. They showed that stock status has been declining in recent years and that current *B* is around *B/B*_MSY_, with the decline being predicted both by individual and ensembled COMs as well as by stakeholders. Our study demonstrates an approach for the data-limited stock assessment of mixed-use fisheries, and complements other recent studies that have shown how marine stock assessment methods can be used in small-scale inland and other small-scale fisheries in transitional waters, such as coastal lagoons (Fitzgerald et al., 2018; Shephard et al., in press). Our work also demonstrates that recreational removals are important to be considered in stock assessments where recreational angling makes up a relevant share of total removals. In our case, neglecting recreational fishing removals lead to an assessment result that indicated a strongly overfished stock due to a long-term decreasing trend in commercial landings. In contrast, by incorporating recreational removals, the total removals time-series trend showed long-term stability of removals up until a recent spike and a subsequent decline, resulting in an assessment that estimates a fully exploited stock status.

Our case study results indicate that the Rügen pike stock is currently fully exploited, but may experience first signs of growth overfishing (*F* > *F*_MSY_). Accordingly, current biomass trends are showing a decline. Commercial and recreational fishing mortality are both relevant factors, and removals from recreational fishing are currently outweighing commercial fisher removals, despite recent catch-and-release rates of pike in Rügen by recreational fishers exceeding 60% (Lucas, 2018). Additionally, environmental changes unrelated to fishing may also contribute to this decline. Previous work on recreational use of pike in inland lakes in the USA has revealed ample variation in recreational fishing-induced pike exploitation rates (Pierce et al., 1995), but the current fishing mortality rate estimated for Rügen pike (*F* = 0.18 yr^-1^) is high when compared with those found in lake studies from the USA (Pierce et al., 1995). Specifically for Germany, roughly 20% of lentic pike stocks have been found to experience fishing mortality rates that are larger than *F*_MSY_ (Arlinghaus et al., 2018), and the Rügen stock thus compares with intensively fished lake stocks.

We assessed the status of the Rügen pike stock using MSY-derived metrics. However, when it comes to mixed-use fisheries, a core question is whether MSY represents the optimal measure for defining stock status (Arlinghaus et al., 2019). For commercial fishers, it has often been suggested that maximum economic yield (MEY) rather than MSY would be a more suitable measure for defining stock status, for two main reasons. Firstly, a stock that is fished at a level that returns MEY instead of MSY would be more desirable from most commercial fishers’ point of view (Norman-López & Pascoe, 2011), whilst secondly the stock would also have a higher overall biomass (Grafton et al., 2007). For recreational fishers, measures of optimal fish status relate more to individual fish size and abundance, rather than maximum biomass yield, and the benefits of a fish stock to recreational fishers are thus usually maximized at lower fishing mortality rates than those that produce MSY (Radomski et al., 2001; Ahrens et al., 2020). For instance, size truncation reduces the satisfaction of those recreational fishers that prefer large pike in the catch (Arlinghaus et al., 2014; Beardmore et al., 2015). Current biomass trends of the Rügen pike are negative and around *B*_MSY_, meaning the stock status is size-overfished and thus may be far from optimal from a recreational fishing point-of-view. In this study we have used MSY as a reference for defining stock status due to its widespread acceptance in fisheries literature as a reference by which stock status should be compared, but acknowledge that this is likely an unsuitable measure of stock status particularly for recreational fishers. We therefore encourage the use of alternative measures by which stock status, and more generally fishery quality, can be quantified in mixed-use fisheries.

The models we have used to derive biomass trends of the Rügen pike stock assumed a constant natural environment. However, this has not been the case in the waters around Rügen. Nutrient load greatly increased from the 1950s to the 1980s (Munkes, 2005), after which it has been steadily declining again (LUNG, 2013). Furthermore, submerged macrophyte coverage has greatly deceased in the Greifswalder lagoon and in the Darss-Zingster lagoons (Kanstinger et al., 2018; Pankow & Wasmund, 1994). Although submerged macrophyte coverage has remained roughly constant over time in the Westrügen lagoons when compared with 1932, it has changed in species composition since then (Blindow et al., 2016). These environmental changes could have affected the productivity of the pike stock, thereby changing the relationship between population abundance and productivity (Vert-Pre et al., 2013), and thus potentially impacting the values of both *B* and *B*_MSY_ over the years (Rose, 2004; Jensen, 2005). In the Darss-Zingster lagoons for instance, increased eutrophication has been thought to be responsible for a decline in pike in favour of pikeperch in the late 1960s (Winkler, 2002). Thus, unaccounted environmental or habitat-driven changes in stock productivity that happened throughout a time-series of landings generally affect stock assessment outcomes and create relevant uncertainty in assessment results (Brown et al., 2019). However, more complex and data-rich age-structured assessment models face a similar issue.

Multiple different COMs have been developed over the years to estimate stock status under data-limited conditions. To be able to run with limited data, these COMs make a variety of simplifying assumptions, increasing the chances for bias and uncertainty of their estimate (Rosenberg et al., 2014; Free et al., 2020). Therefore, using only a single COM for assessing stock status increases the risk of producing a flawed or biased status estimate. This is supported by the results of our case study, which showed a large spread in individual COM predictions. Thus, using only a single COM to estimate stock status should be avoided. Instead, the use of multiple different models in an ensemble approach, with the final estimated value either being the average of all models or the product of some weighting procedure, is expected to increase the robustness of the result (Bates & Granger, 1969).

Although a simple model averaging approach could improve estimates of stock status, it will not incorporate that some models perform better or worse under certain conditions. A superensemble model, on the other hand, allows for exploiting the covariance between individual COM predictions (Anderson et al., 2017), allowing for a better accounting of individual model biases. In our case study, we applied three different types of superensemble models and found their estimates of recent biomass status were relatively similar, increasing our confidence in their results. However, Free et al. (2020) found that superensemble models of COMs generally produce a negatively-biased estimate of stock status for lightly-exploited stocks. Thus, it is possible that the pike stock in our case study is actually in a better shape than suggested by the superensemble models.

Furthermore, the predominant reliance of COMs on landings time-series means that it is important that these data are reliable and of high quality. For many mixed-use fisheries this is rarely the case, with data on recreational removals often missing or being incomplete (Arlinghaus et al., 2019). Commercial landings statistics may also suffer from illegal and unreported catch (Agnew et al., 2009). In our case study, we reconstructed a time-series of recreational removals using two local scientific studies as anchor points, and using various other forms of time-series data to interpolate between and extrapolate from these points. Such reconstructions should be paired with rigorous sensitivity analyses. Even though we tested the sensitivity of the model results to various alternative reconstruction approaches and found limited impact, we cannot exclude the possibility that our reconstruction of recreational removals contained uncertainty. Nevertheless, our results showed that it is advisable to reconstruct recreational fishing removals, even when uncertain, instead of solely relying on commercial landings data. Thus, in the absence of sufficient data to perform more sophisticated assessments that incorporate age data, COMs may provide approximations of stock status, as long as uncertainties are recognized and explored. We tested several sensitivities and found our results to be largely robust, and additionally mirrored by stakeholder perceptions.

There is still an active debate among fisheries scientists on whether catch-only methods should be used at all for assessing fish stock status. Some argue that catch data represents the only data available for many data-limited fish stocks, and that using catch-only methods provides the only option of getting an indication of the status of those stocks, even if they are less precise than data-rich stock assessments (Froese et al., 2012; Pauly et al., 2013). Others argue that it is better not to use catch-only methods when they may be wrong, and that instead the focus should be on collecting and including additional data (Branch et al., 2011; FAO, 2019). Explicitly accounting for the higher levels of uncertainty of data-limited methods by including precautionary management measures or buffers may resolve some of this debate (Dowling et al., 2019). For instance, Walsh et al. (2018) found that using superensembles of catch-only models to inform fisheries management could reduce the risk of overfishing, but only when combined with a precautionary harvest control rule, which in turn might result in poor yields. Thus, the usage of COMs and superensembles in the management of mixed-use fisheries should be combined with a precautionary approach, and whenever possible additional data should be collected to allow for more data-rich assessments. For instance, we added survey data from stakeholders to increase our confidence in the assessment outcomes. Although stakeholder perspectives present their own biases (O’Donnell et al., 2010), the alignment of stakeholder perceptions and COM predictions support the conclusion that pike biomass has declined in recent years.

## Conclusion

We have used a combination of individual and ensembled COMs and stakeholder surveys to assess the status of a data-poor coastal pike stock that is exploited by both commercial and recreational fishers. We conclude that the pike stock is fully exploited, currently declining, and may be experiencing growth overfishing. Thus, reductions in fishing mortality may be advisable. Our study has shown the benefits of using multiple different models and including stakeholder surveys when assessing stock status through data-limited methods, and has also demonstrated the importance of including recreational removals when assessing the status of a stock that is co-exploited by commercial and recreational fisheries.

## Acknowledgements

Funding was received through the European Maritime Fisheries Fund (EMFF) of the EU and the State of Mecklenburg-Vorpommern (Germany) (grant MV-I.18-LM-004, B 730117000069) within the Boddenhecht project. We thank Chris Free, Olaf Jensen, Dan Ovando, and Robert Ahrens for discussion, and furthermore thank Chris Free for the sharing of model code and a detailed review of the manuscript draft. There is no conflict of interest.

