## Appendix A for "Data-poor stock assessment of fish stocks co-exploited by commercial and recreational fisheries: applications to pike (*Esox lucius*) in the western Baltic Sea"

Appendix A – Recreational removals reconstruction

Recreational removals of pike from the lagoons around the German Baltic island of Rügen were reconstructed for the period 1955-2018, using two regional angling studies, commercial catch data, and various proxies for commercial and recreational effort.

*Commercial removals data*

Annual commercial landings from 1976 to 2018 were obtained from the Landesamt für Landwirtschaft, Lebensmittelsicherheit und Fischerei (LALLF), which is the state office for agriculture, food safety and fisheries of the German state of Mecklenburg-Vorpommern (M-V). Furthermore, data of annual commercial pike landings from 1969-1975 and from 1955-1968 were extracted from Winkler (1991) and and from records generated from annual official fisheries reports published by the Institut für Hochseefischerei und Fischverarbeitung Rostock of the former GDR, respectively. Thus, a time-series of annual commercial pike landings in the Rügen area is available for the years 1955-2018 (Figure A1).

*Commercial effort data*

No effort data is available for individual fishers. Instead, as a proxy for fishing effort in a given year, the number of active fishing vessels in that year is used. This data is extracted from the European Union Fleet Register as the number of fishing vessels with an active fishing license that are registered to a port within one of the relevant districts around Rügen. Furthermore, all fishing vessels with a total length of 20 meters or more are discarded from the data, as these are assumed to not be fishing in coastal waters but in the deeper waters of the Baltic Sea. The EU Fleet Register also provides some possible alternative measures of fishing effort, e.g. vessel length, tonnage, and power. However, we expect that these are better effort proxies for active fishing gear such as trawl nets, and are less suitable as an effort proxy for passive gear such as the gill nets and fyke nets in which pike are caught. We therefore use the number of active fishing vessels in the region as a proxy for fishing effort.

The EU fleet register has data on German vessel numbers from 1991 up to 2018. However, it appears that not all vessels that were active in 1991 are listed in the register, as the number of registered fishing vessels increased from 874 in 1991 to 1080 in 1992, after which it almost continually declines. This could be related to the reunification of Eastern and Western Germany that took place around this time. Therefore, we discard the 1991 vessel numbers, giving us a proxy for fishing effort in the region from 1992 up to 2018 (Figure A2).

For the years 1955-1991, no data is available on the number of commercial fishing vessels. However, to be able to reconstruct recreational catches for this time period, some estimate of commercial effort is needed during this time. We therefore make the assumption that commercial effort was constant in this time period (a rather unrealistic assumption, but one that must be made if we want to make some estimate of recreational catches during this period). Thus, it is assumed that commercial effort in 1955-1991 was the same as in 1992. (Figure A2).

*Recreational effort data*

We have no direct measure of recreational effort in the Rügen area for the years 1955-2018. Instead, we used several different proxies for recreational effort to reconstruct a recreational effort time-series.

The first proxy we used was the number of angling license stamps (“Fischereiabgamarke” in German) to get an indication of the total number of resident anglers in M-V. To have a valid fishing license, an M-V resident must buy one of these stamps, and attach it to their fishing license. The stamp is valid for one calendar year, and must thus be renewed each year to keep a valid fishing license. Data on the annual number of angling license stamps was obtained from LALLF.

The second proxy we used was the number of coastal angling licenses issued by the state of M-V, which are available from 1991 to 2018 (Figure A3). These licenses allow the bearer to fish in all coastal waters of M-V, and must be bought by both resident and tourist anglers if they want to be able to legally fish in M-V coastal waters. Several different versions of the coastal fishing license are issued: one that is valid for a year, one that is valid for a week, and one that is valid for a day. Furthermore, in 2004 a youth license was introduced that is valid for a year and can only be bought by anglers under 18 years of age, and in 2010 a disabled license was introduced that is valid for a year and can only be bought by anglers with a disability.

The third proxy we used was membership data of the Deutsche Anglerverband (DAV), which was the national recreational angling association in the GDR. DAV membership data was available for the years 1954 (the year of its founding), 1960, 1970, 1980, and 1990 (VDFF, 1998). Membership data for the missing years between 1954 and 1990 was estimated by linearly interpolating between known data points (Figure A4).

The fourth proxy we used were two telephone-diary studies performed in 2006-2007 (Dorow & Arlinghaus, 2011) and 2014-2015 (Weltersbach et al., in press; summarized in Lucas, 2018). The Dorow & Arlinghaus (2011) study provided an estimate of the number of resident M-V anglers that fished in the Rügen region for the 2006-2007 season relative to the total number of resident M-V anglers, and the average number of angling trips an average resident M-V angler took to the Rügen area. The Strehlow et al. (in press) study provided an estimate of the number of both resident and tourist angling trips that were made in the Rügen region in the 2014-2015 season. The Dorow & Arlinghaus (2011) is assumed to be representative of the 2007 calendar year, and the Weltersbach et al. (in press) study is assumed to be representative of the 2015 calendar year.

*Reconstructing recreational effort*

We used the available data of the above recreational effort proxies to reconstruct a time-series of recreational angling trips taken in the Rügen area from 1955 to 2018.

First, we estimated the number of resident M-V angler trips for 2007. For this, we first calculated the number of angling trips in the Rügen area per resident M-V angler by multiplying the estimate of the number of resident M-V anglers that fished in the Rügen region for the 2006-2007 season relative to the total number of resident M-V anglers with the average number of angling trips an average resident M-V angler took to the Rügen area. Multiplying with the number of angling license stamps for 2007 gave an estimate of the number of resident angler trips in the Rügen area in 2007, assuming that the number of angling license stamps reflects the total number of residential M-V anglers.

Next, we constructed a time-series of resident angler trips per annual coastal angling license for both 2007 and 2015. We first divided the estimated number of resident angler trips of both 2007 and 2015 that year’s number of annual coastal angling licenses (including the youth and disabled license), assuming that only resident anglers apply for a coastal angling license that is valid for a full year. This gave an estimate of resident angler trips per coastal license for 2007 and 2015. We then estimated a time-series of resident angler trips per annual coastal angling license for the years 1991 to 2018 by linearly interpolating between 2007 and 2015, and by assuming that all years prior to 2007 have the 2007 value and all years after 2015 have the 2015 value.

We then reconstructed a 1991-2018 time-series of number of resident angler trips in the Rügen region by multiplying each year’s value of resident angler trips per annual coastal angling license with the number of annual coastal angling licenses sold in that year (including the youth and disabled licenses).

Next, we reconstructed the number of tourist angler (not from M-V) trips in the Rügen region. We first divided the number of tourist angler trips taken in 2015 by that year’s number of short-term (valid for a day and a week) coastal angling licenses (assuming that only tourists buy the short-term coastal angling licenses). Assuming a constant ratio of tourist angler trips in the Rügen area to short-term coastal angling licenses, we multiplied the 2015 ratio with each year’s number of short-term coastal angling licenses, thereby reconstructing the number of tourist angler trips in the Rügen region for the years 1991-2018.

Summing up the number of resident and tourist angler trips in the Rügen area gave an estimate of total angler trips in the Rügen area for the years 1991-2018. We then reconstructed the number of angler trips in 1955-1990 by using the DAV membership data. First, we linearly extrapolated the estimated number of Rügen angler trips in 1990 from the number of trips in 1991-2000, which shows a near-perfect linear increase (Figure A5). For 1990, we subsequently calculated the number of Rügen angler trips per DAV member. The number of Rügen angler trips from 1954 to 1990 is then reconstructed by assuming that the estimated fraction of Rügen angler trips per DAV member is constant for all years of DAV membership data (Figure A5).

We have now reconstructed a time-series of recreational angling trips in the Rügen area for 1955-2018. Using data on recreational removals, we can use this to reconstruct a time-series of recreational removals.

*Recreational removal data*

Contrary to commercial catch data, there is little information available on recreational removals of pike around Rügen, as this is not actively monitored in M-V. The only data available on recreational angling catches of pike in the region are from the two telephone-diary studies performed in 2006-2007 (Dorow & Arlinghaus, 2011) and 2014-2015 (Strehlow et al., in press; summarized in Lucas, 2018). Dorow & Arlinghaus (2011) provide information on per-trip average catch, harvest, and release rate of pike (in numbers) by resident anglers. Strehlow et al. (in press) provide information on per-trip average catch, harvest, and release rate of pike (in numbers and biomass) by both resident and tourist anglers. Furthermore, we calculated the average size of a caught pike in 2015 by dividing the 2015 per-unit-biomass catch rate by the per-number catch rate. The above data is used to estimate recreational pike removals in 2007 and 2015, and are subsequently used to reconstruct recreational pike removals for the remaining years between 1955 and 2018.

*Reconstructing recreational removals*

We estimated resident angler catch, harvest, and release numbers in 2007 by multiplying the 2007 resident angler per-number catch, harvest, and release rate with the estimated number of resident angler trips in the Rügen area in 2007. Multiplying with the average size of caught pike in 2015 gave an estimate of 2007 resident angler catch, harvest, and release (assuming that the average size of a captured pike remained the same).

Next, we estimated resident angler catch, harvest, and release in 2015 in terms of biomass by multiplying the 2015 resident angler per-unit-biomass catch, harvest, and release rate with the estimated number of resident angler trips in the Rügen area in 2015. Similarly, we estimated tourist angler catch, harvest, and release in 2015 in terms of biomass by multiplying the 2015 tourist angler per-unit-biomass catch, harvest, and release rate with the estimated number of tourist angler trips in the Rügen area in 2015.

Then, we reconstructed a recreational catch time-series for 1991-2018 by estimating a recreational catch-per-unit-effort (CPUE) time-series for both resident and tourist catches. We did this by calculating a CPUE constant of proportionality between commercial and recreational catches according to:

|  | ${C_{R,t}}/{E_{R,t}}=c_{\mathrm{CPUE}}{C_{C,t}}/{E_{C,t}}$ | A1 |
| --- | --- | --- |

Where $c_{\mathrm{CPUE}}$ is the CPUE constant of proportionality, $C_{R,t}$ and $C_{C,t}$ are the recreational angler and commercial catches at time $t$ respectively, and $E_{R,t}$ and $E_{C,t}$ are the measures for recreational angler and commercial effort respectively (number of trips and number of registered fishing vessels). For resident angler catches, $c_{\mathrm{CPUE}}$ was calculated for both 2007 and 2015, and the average was taken. For tourist catches, $c_{\mathrm{CPUE}}$ was calculated with only the 2015 data. We then calculated a CPUE time-series for both resident and tourist angler catches from 1991 to 2018 by multiplying the respective $c_{\mathrm{CPUE}}$ value with each year’s value of commercial CPUE. Lastly, multiplying each year’s resident and tourist CPUE with that year’s respective number of resident and tourist angling trips gave a time-series of resident and tourist catches.

We have now reconstructed a time-series of resident and tourist angler catches for 1991-2018. However, not all fish that are caught by anglers are harvested. Therefore, to reconstruct a time-series of actual angler removals, we first accounted for release rates. To account for changes over time in the release rate of resident anglers, we reconstructed a time-series of resident angler release rate for 1991-2018. For this, we used the resident angler release rates of 2007 and 2015. We linearly interpolated for the years between 2007 and 2015, while assuming that release rate for all years before 2007 was equal to that of 2007, and assuming that release rate for all years after 2015 was equal to that of 2015. For tourist release rates we only have information from 2015, so we assumed that tourist angler release rate was constant for all years, and equal to the 2015 rate.

By assuming a pike release mortality $\mu_{R}$ of 7.8% (Hühn & Arlinghaus, 2011), total pike removals $T$ by resident and tourist anglers for 1991-2018 can now be reconstructed according to:

|  | $T_{i,t}=C_{i,t}\left( 1-r_{i,t} \right)\mu_{R}r_{i,t}$ | A2 |
| --- | --- | --- |

where $C_{i,t}$ is catch in year $t$ , $r_{i,t}$ is release rate in year $t$, and $i$ denotes either resident or tourist anglers. By summing resident and tourist removals, we obtained a time-series of total angling removals in the Rügen area for 1991-2018.

No information is available on separate resident and tourist angler effort before 1991, or changes in their release rate. Therefore, we reconstructed total angler removals for the years 1955-1990 by calculating a CPUE constant of proportionality $c_{\mathrm{CPUE}}$according to Equation A1 with 1991 data, thereby using total 1991 angler removals instead of catches as $C_{R,t}$. We multiplied this value of $c_{\mathrm{CPUE}}$ with each year’s commercial CPUE for 1955-1990, thereby obtaining a time-series or recreational CPUE for 1955-1990. Multiplying each year’s value of recreational CPUE with recreational effort yielded a time-series of total recreational removals for 1955-1990 (Figure 2, main text).

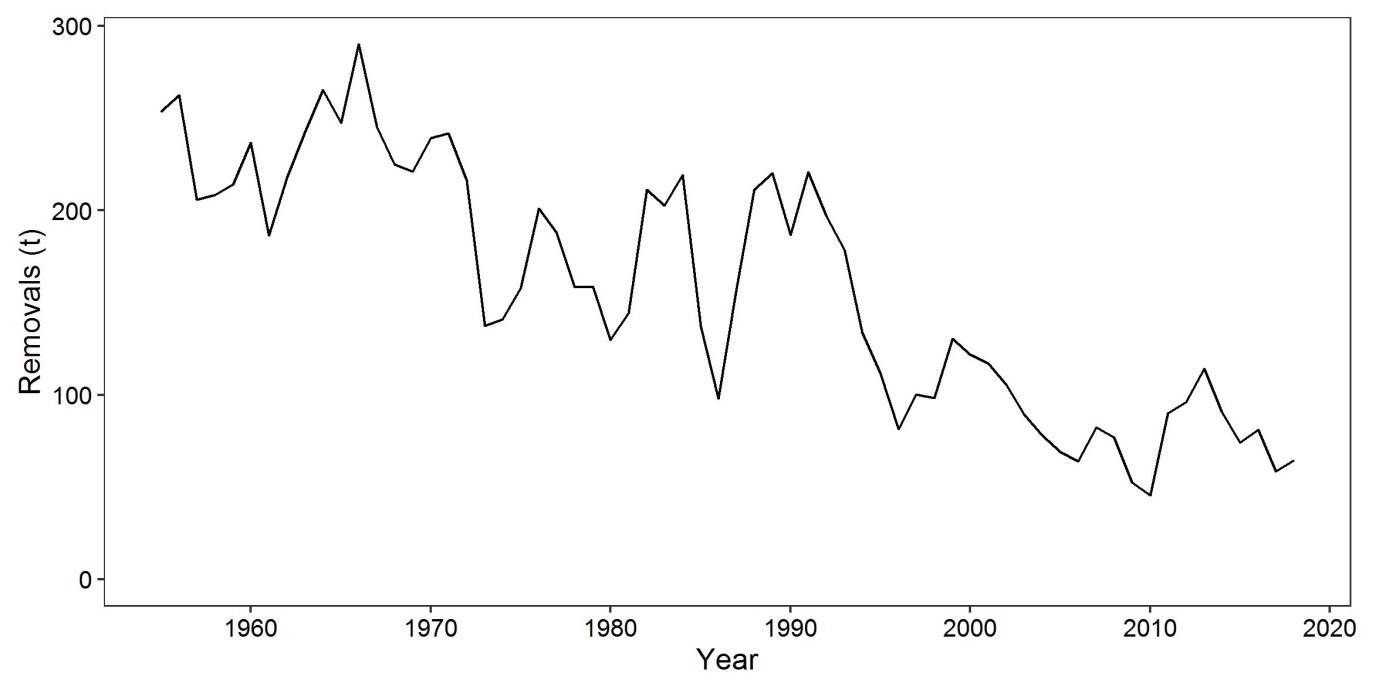

Figure A1: Commercial fisheries landings of pike in the Rügen area over time. 1955-1975 data came from Winkler (1991) and further supplemented by records generated from annual official fisheries reports published by the Institut für Hochseefischerei und Fischverarbeitung Rostock of the former GDR. 1976-2018 data came from Mecklenburg-Vorpommern’s Landesamt für Landwirtschaft, Lebensmittelsicherheit und Fischerei (LALLF).

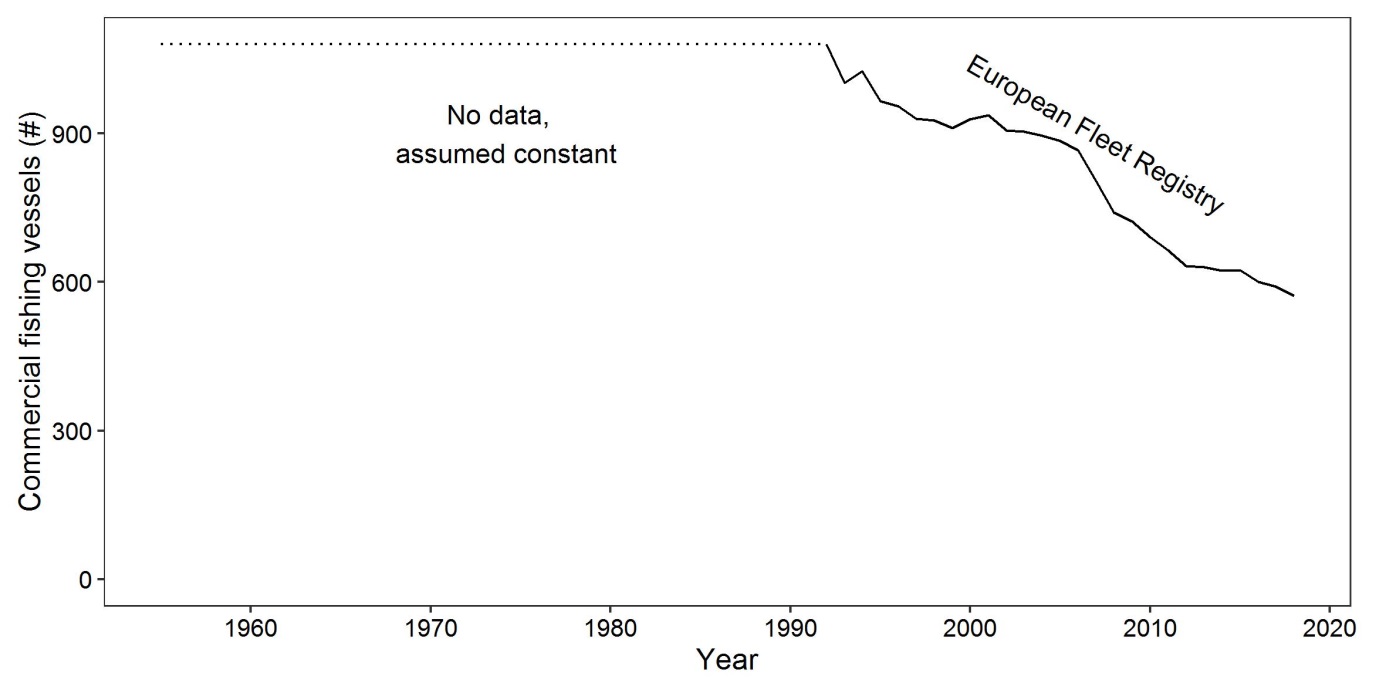

Figure A2: Number of commercial fishing vessels registered to a port in the Rügen area over time, used as a proxy for commercial fishing effort. Vessel numbers from 1992 to 2018 were taken from the European Fleet Registry (solid), while the number of commercial fishing vessels for 1955-1991 were assumed to be the same as that in 1992 (dashed).

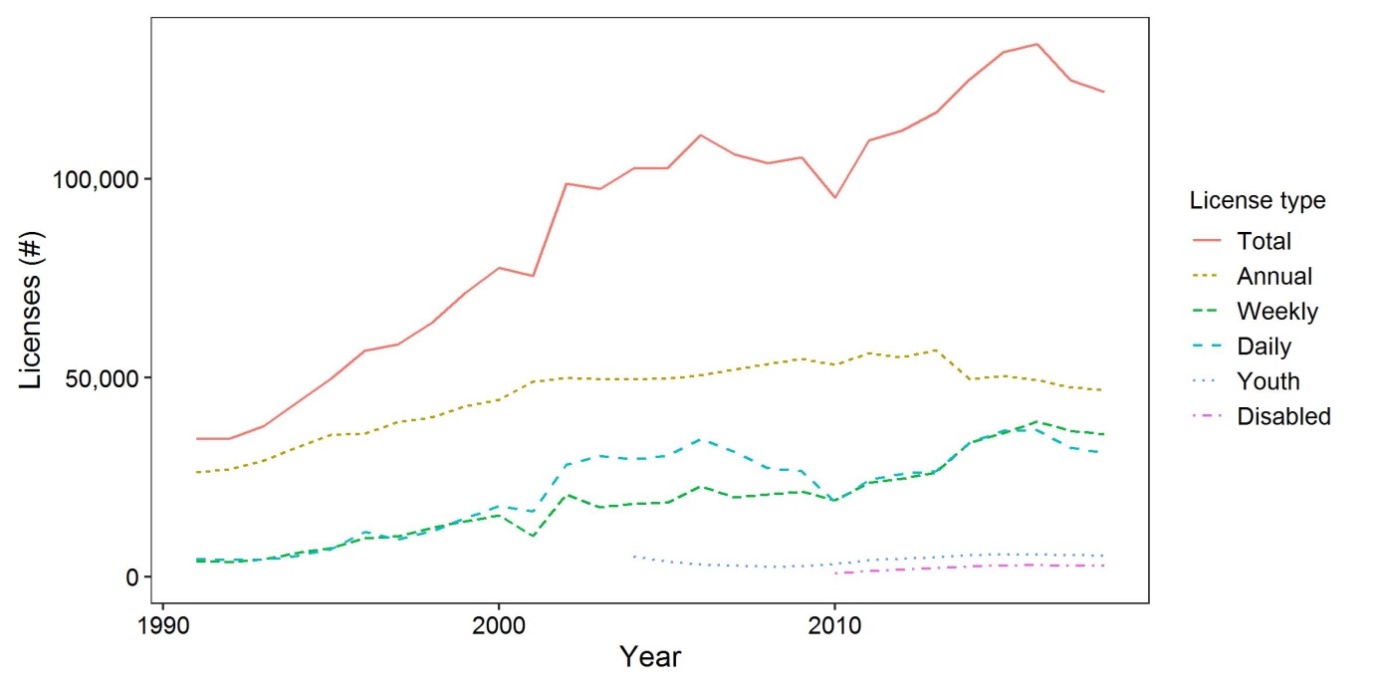

Figure A3: Number of coastal fishing licenses purchased in the state of M-V over time. Data obtained from LALLF.

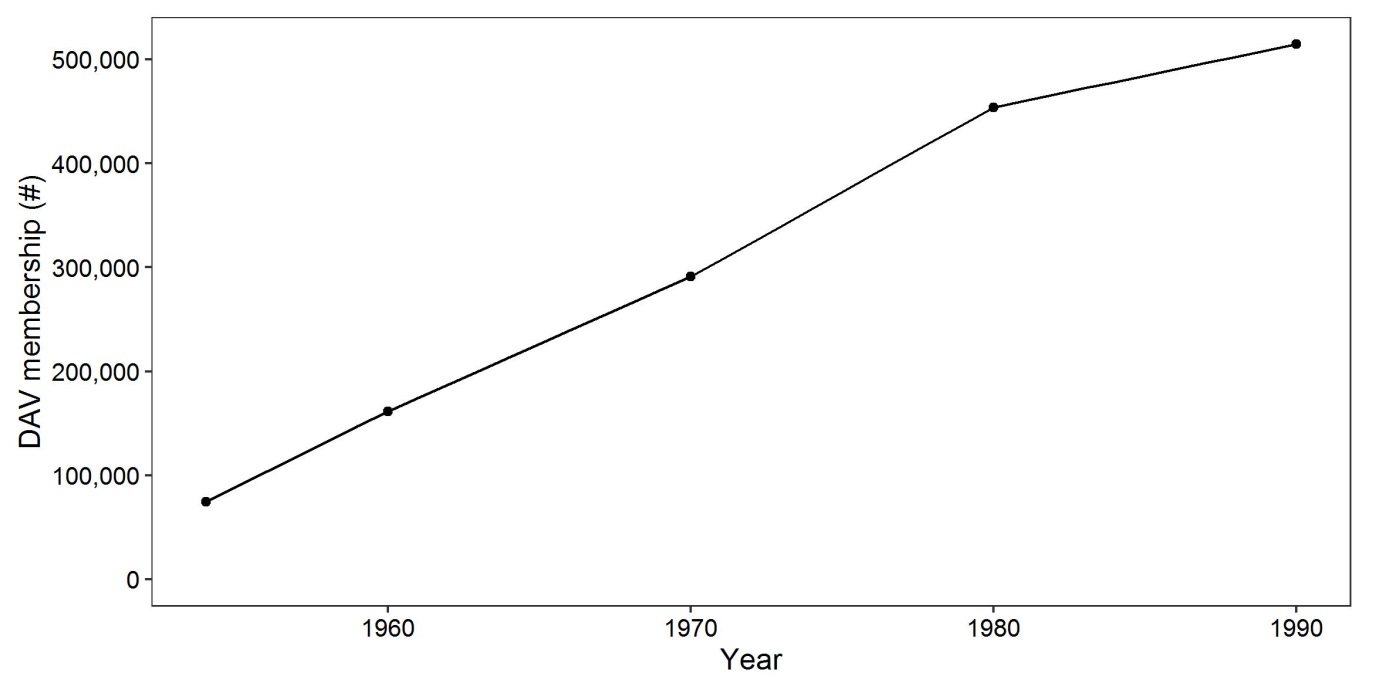

Figure A4: Membership numbers of the Deutscher Anglerverband (DAV) over time, which was the national angling association of the GDR. The points show known membership numbers, given by VDFF (1998). The intermediate years (solid line) have been estimated through linear interpolation.

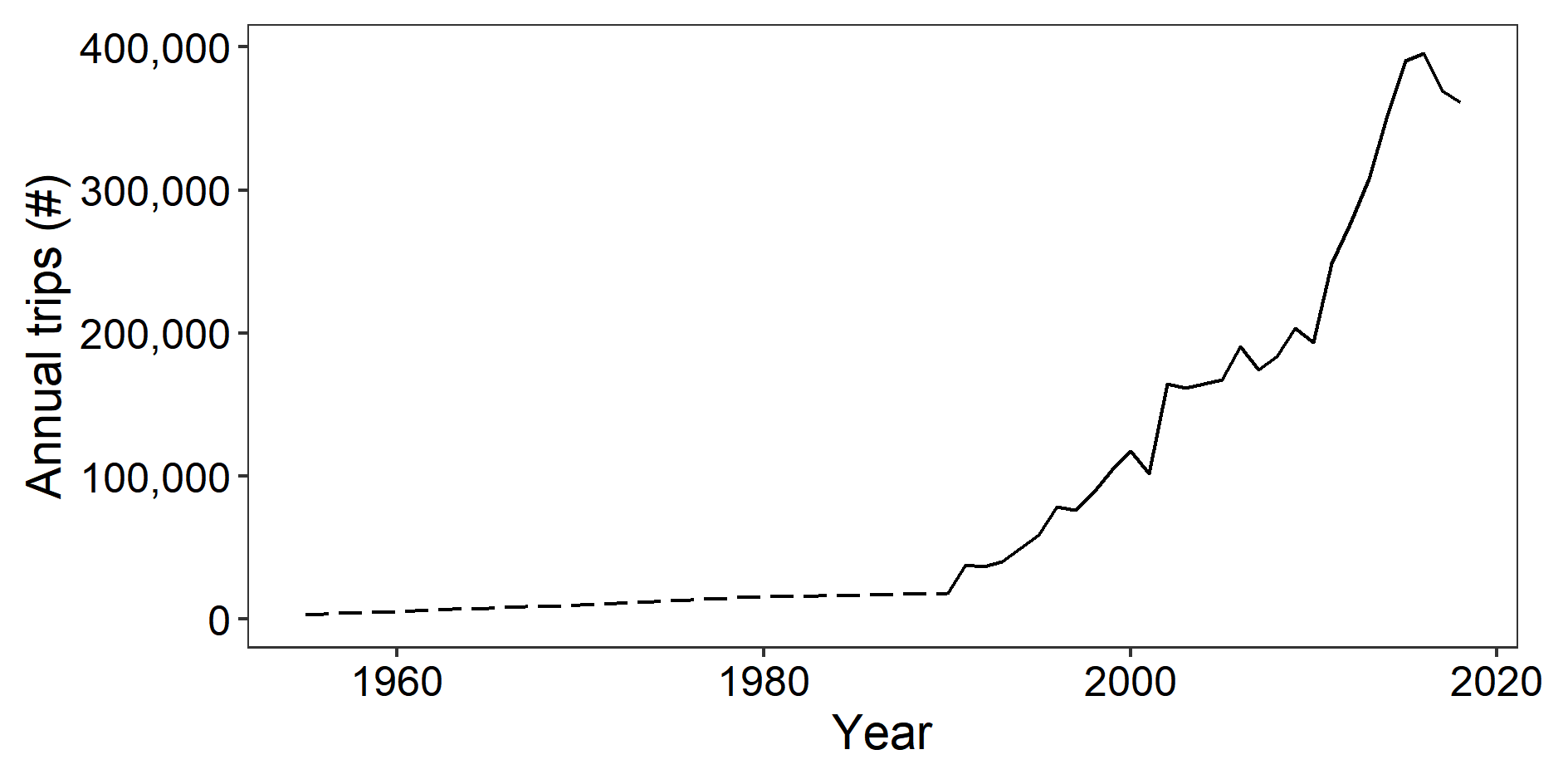

Figure A5: Reconstructed number of annual trips taken to the Rügen area by recreational anglers over time. The dashed line has been reconstructed based on DAV membership data (Figure A4), and the solid line has been reconstructed based on coastal fishing license data from M-V (Figure A3).
