## Appendix B for "Data-poor stock assessment of fish stocks co-exploited by commercial and recreational fisheries: applications to pike (*Esox lucius*) in the western Baltic Sea"

Appendix B – Description of catch-only models (COMs)

Catch-MSY uses a catch time-series, estimates of relative stock size in the first and final year of that time-series, and prior ranges for population production parameters $r$ (intrinsic population growth rate) and $k$ (population carrying capacity) as model input. With this data, a Schaefer production model is run for each $r$-$k$ pair, and all non-viable $r$-$k$ pairs are discarded (those that result in unrealistic biomass trajectories). An estimate of MSY is then derived from all viable runs (Martell & Froese, 2013). We used the modifications introduced by Rosenberg *et al.* (2014) to find $B/{B_{\mathrm{MSY}}}$ as the median of the $B/{B_{\mathrm{MSY}}}$ values of all viable runs. As inputs to Catch-MSY, we assumed stock size in the starting year to be between 20% and 80% of unexploited biomass (a broad range due to lack of knowledge), stock size in the final year to be between 20% and 70% of unexploited biomass (assuming a lower upper range due to technological improvements in fishing over time), $r$ to be between 0.05 and 0.5 yr^-1^ (following Martell & Froese, 2013, in translating a “low” resilience as given by Froese & Pauly, (2019), to an r-range), and $k$ to be between the highest observed catch and 50 times the highest observed catch ($k$ is unlikely to be smaller than the highest observed catch, and under a targeted fishery also unlikely to be many orders of magnitude greater). Aside from $B/{B_{\mathrm{MSY}}}$, Catch-MSY also returns an estimate of MSY, $B_{\mathrm{MSY}}$, and time-series $B$. We calculated time-series $F$ as catch divided by $B$, and $F_{\mathrm{MSY}}$ as MSY divided by $B_{\mathrm{MSY}}$. In general, Catch-MSY’s $B/{B_{\mathrm{MSY}}}$ estimates were found to be bimodal: giving either a low or high estimate of $B/{B_{\mathrm{MSY}}}$ and no intermediate estimate (Free et al., 2020).

CMSY functions similarly to Catch-MSY, but differs in how it derives MSY and its associated reference points from the viable $r$-$k$ pairs. CMSY looks for the most likely $r$-$k$ pair in the peak of the triangle of all viable $r$-$k$ pairs, as $r$ represents the *maximum* population growth rate and that peak is where the largest values of $r$ can be found (Froese et al., 2017). Furthermore, CMSY introduces a linear decline of surplus production if biomass falls below $0.25k$, to account for production models tending to overestimate surplus production at low stock sizes (Froese et al., 2017). As inputs to CMSY, we assumed stock size in the starting year to be between 20% and 80% of unexploited biomass, stock size in the final year to be between 20% and 80% of unexploited biomass, $r$ to be between 0.05 and 0.5 yr^-1^, and $k$ to be between the highest observed catch and 50 times the highest observed catch. Aside from $B/{B_{\mathrm{MSY}}}$, CMSY also returns an estimate of MSY, $B_{\mathrm{MSY}}$, $F_{\mathrm{MSY}}$, time-series $B$, and time-series $F$. In general, CMSY’s $B/{B_{\mathrm{MSY}}}$ estimates were found to be negatively-biased (on average estimating a $B/{B_{\mathrm{MSY}}}$ value that is lower than the true state), and becoming more accurate with greater fishing pressure (Free et al., 2020).

COM-SIR (catch-only model with sampling-importance resampling) uses a catch time-series and prior ranges for population production parameters $r$ (intrinsic biomass growth rate), $k$ (biomass carrying capacity), $x$ (intrinsic rate of effort increase), and $a$ (fraction of $k$ at which there is bioeconomic equilibrium) as model input to reconstruct historical biomass and effort, by assuming that biomass follows a Schaefer production model and that effort moves toward the bioeconomic equilibrium via a logistic function (Vasconcellos & Cochrane, 2005). For this, parameters are estimated using the Bayesian sampling importance resampling (SIR) algorithm (McAllister et al., 1994). As input to COM-SIR, we assumed $r$ to be between 0.05 and 0.5 yr^-1^, $k$ to be between the maximum observed catch value and 50 times the maximum observed catch value, $x$ to be between 1E-6 and 1 (Rosenberg et al., 2014), and $a$ to be between 0 and 1 (Rosenberg et al., 2014). Aside from $B/{B_{\mathrm{MSY}}}$, COM-SIR returns estimates of time-series $B$ and time-series $F$. MSY, $B_{\mathrm{MSY}}$, and $F_{\mathrm{MSY}}$ were estimated from the median values of the posterior distributions of $r$ and $k$: MSY was estimated as $rk/4$, $B_{\mathrm{MSY}}$ as $k/2$, and $F_{\mathrm{MSY}}$ as $r/2$. In general, COM-SIR’s $B/{B_{\mathrm{MSY}}}$ estimates were found to be positively-biased, especially when the stock is overexploited (Free et al., 2020).

SSCOM (state-space catch only model) uses a time-series of catch and an estimate of the intrinsic biomass growth rate $r$ as model input. By assuming that effort follows predictable dynamics (moving net fisheries profit towards 0) and assuming that biomass follows a Schaefer production model, SSCOM recreates population biomass and effort from the catch time-series by making use of Markov chain Monte Carlo sampling (Thorson et al., 2013). As input to SSCOM, we assumed $r$ to be between 0.05 and 0.5 yr^-1^. In general, SSCOM’s $B/{B_{\mathrm{MSY}}}$ estimates were found to be positively-biased, especially when the stock is overexploited (Free et al., 2020).

mPRM (modified panel-regression model) uses a time-series of catch and broad life-history information as model input (Rosenberg et al., 2014), and is a modified version of the PRM method described in Costello et al. (2012). First, mPRM takes data from the RAM legacy stock assessment database (Ricard et al. (2012) to fit a regression model to catch data and a species fixed effect, using assessed $B/{B_{\mathrm{MSY}}}$ as the response variable. Then, by inputting the catch data and species group of the unassessed stock, the regression coefficients are used to estimate stock status of the unassessed stock. Here, we categorize Baltic pike under the species category "Miscellaneous coastal fishes". In general, mPRM’s $B/{B_{\mathrm{MSY}}}$ estimates were found to become more accurate with greater fishing pressure (Free et al., 2020).

zBRT (Zhou boosted regression trees) uses a time-series of catch to estimate stock saturation (biomass relative to unfished biomass, $B/k$) by training a boosted regression tree model on the catch data of data-rich assessed stocks in the RAM Legacy database, and subsequently applying this trained model to the data-poor stock in question (Zhou et al., 2017). 56 different predictor variables are considered when building the model, which are derived from the catch data (e.g. the regression coefficients of the scaled catch before and after the maximum catch). $B/{B_{\mathrm{MSY}}}$ is derived from the $B/k$ estimate by assuming that $B_{\mathrm{MSY}}=0.5k$. In general, zBRT was found to provide reasonable estimates of $B/{B_{\mathrm{MSY}}}$ (Free et al., 2020).

OCOM (optimized catch-only model) uses a time-series of catch and a prior for natural mortality $M$ as model input, deriving a prior for $r$ from the prior for $M$ through life history correlations, and assuming that biomass at the start of the catch time series equals $k$ (Zhou et al., 2018). OCOM produces estimates of $B/{B_{\mathrm{MSY}}}$ as well as other population parameters and fisheries reference points by first obtaining an estimate of $B/k$ in the final year of the catch time series by making use of the zBRT model, and then using a Schaefer production model together with an optimization algorithm to find the value of $k$ for which $B/k$ in the final year is closest to that estimated by zBRT (Zhou et al., 2018). Here, we assume that $M$ equals 0.268 yr^-1^ following Ahrens *et al*. (2020). Aside from $B/{B_{\mathrm{MSY}}}$, OCOM also returns an estimate of MSY, $B_{\mathrm{MSY}}$, $F_{\mathrm{MSY}}$, time-series $B$, and time-series $F$. In general, OCOM’s $B/{B_{\mathrm{MSY}}}$ estimates were found to become more accurate with greater fishing pressure, and were found to be bimodal: giving either a low or high estimate of $B/{B_{\mathrm{MSY}}}$ and no intermediate estimate (Free et al., 2020).
