## Appendix C for "Data-poor stock assessment of fish stocks co-exploited by commercial and recreational fisheries: applications to pike (*Esox lucius*) in the western Baltic Sea"

Appendix C – Superensemble model tuning

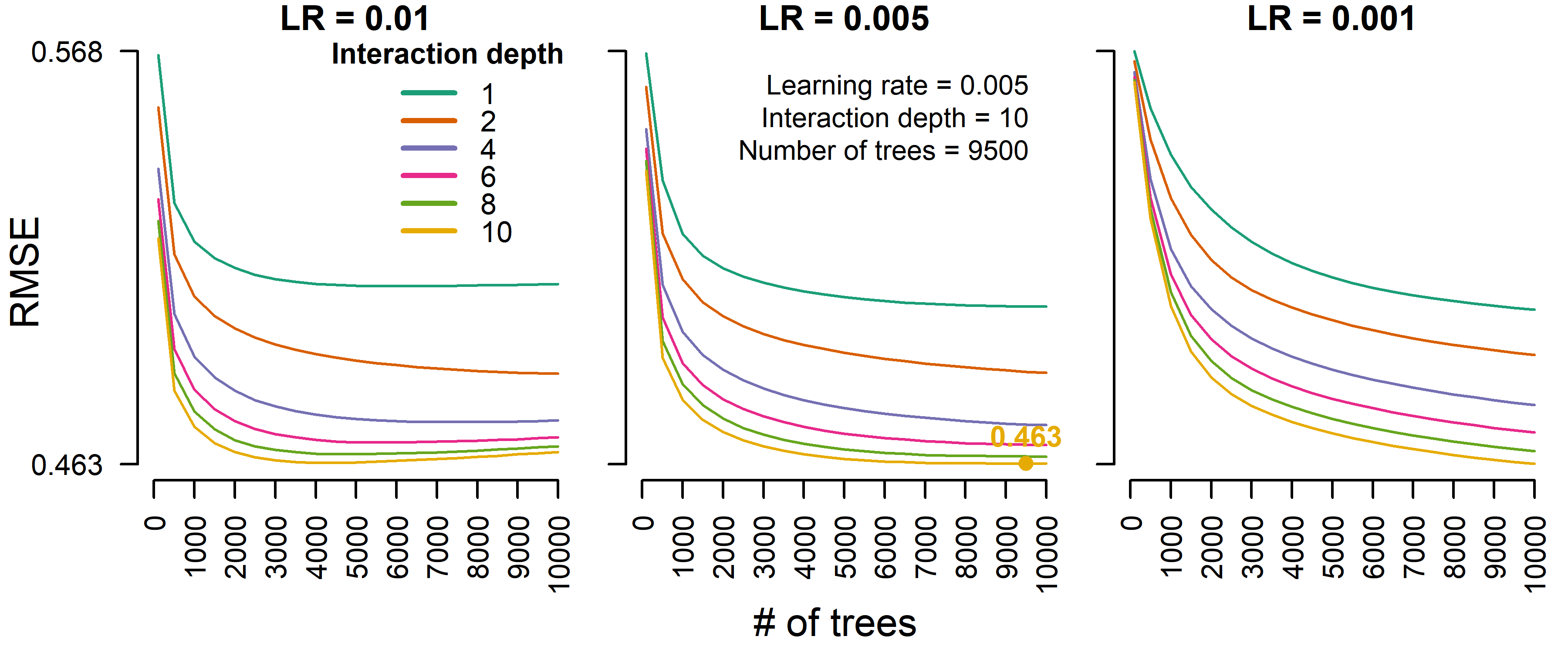

Figure C1: Boosted regression tree model tuning results for mean $\boldsymbol{B}/{\boldsymbol{B}_{\mathbf{MSY}}}$ of the last 5 years of data. The parameter values that resulted in the smallest root-mean-square deviation were chosen for the final boosted regression tree model.

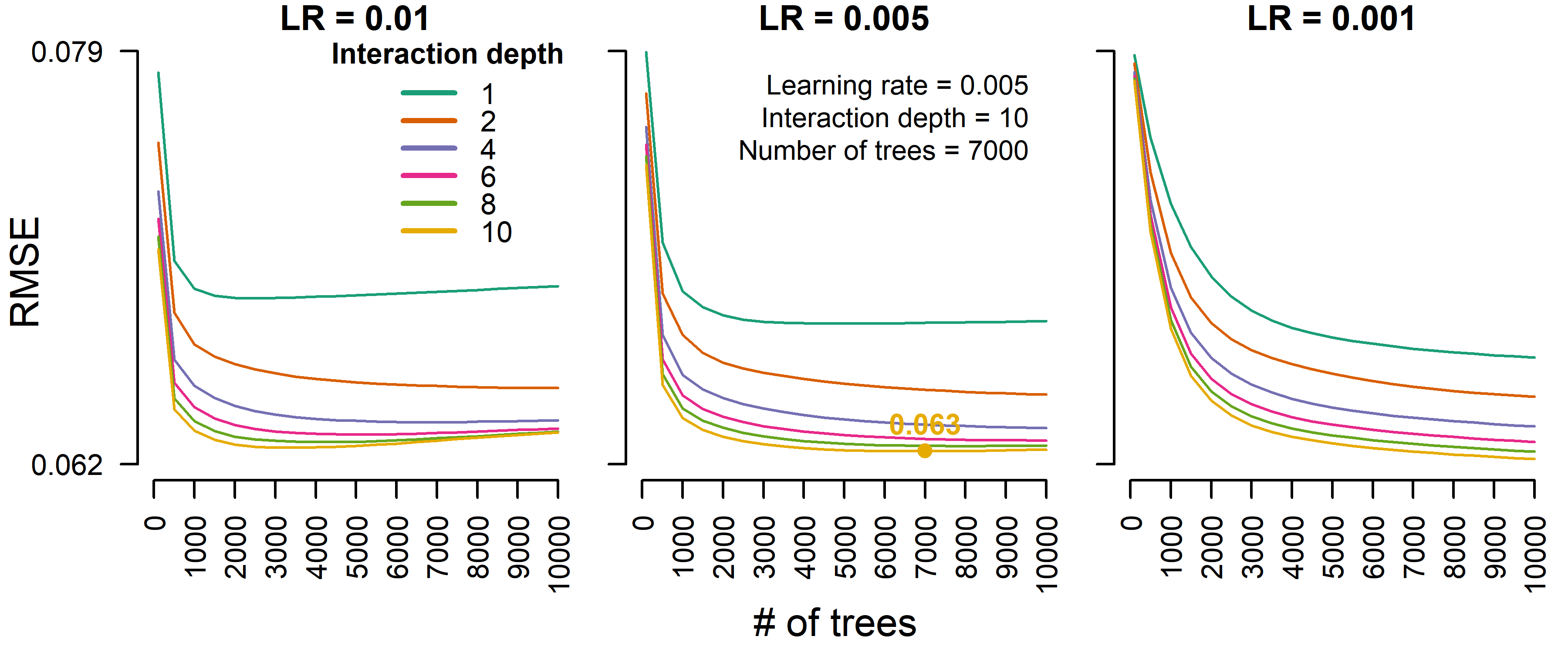

Figure C2: Boosted regression tree model tuning results for $\boldsymbol{B}/{\boldsymbol{B}_{\mathbf{MSY}}}$ slope of the last 5 years of data. The parameter values that resulted in the smallest root-mean-square deviation were chosen for the final boosted regression tree model.

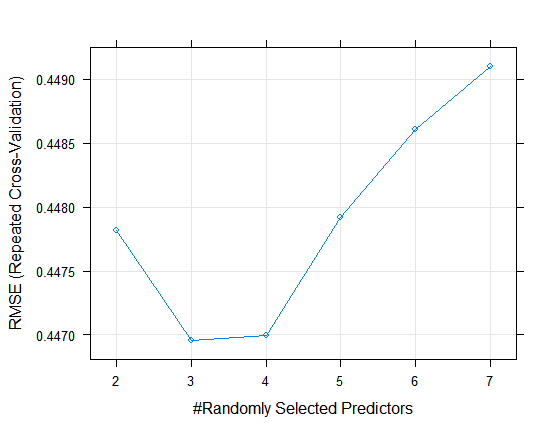

Figure C3: Random forest model tuning results for mean $\boldsymbol{B}/{\boldsymbol{B}_{\mathbf{MSY}}}$ of the last 5 years of data. The parameter value that resulted in the smallest root-mean-square deviation were chosen for the final random forest model.

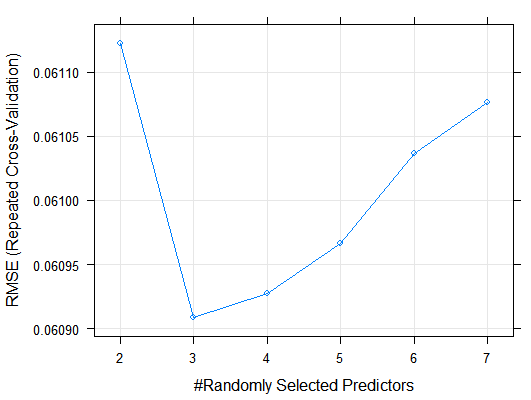

Figure C4: Random forest model tuning results for $\boldsymbol{B}/{\boldsymbol{B}_{\mathbf{MSY}}}$ slope of the last 5 years of data. The parameter value that resulted in the smallest root-mean-square deviation were chosen for the final random forest model.
