## Appendix D for "Data-poor stock assessment of fish stocks co-exploited by commercial and recreational fisheries: applications to pike (*Esox lucius*) in the western Baltic Sea"

Appendix B – Sensitivity analysis

We performed a sensitivity analysis to test the influence of catch-only model (COM) input parameter values and various methods of reconstructing the pike total removals time-series on the results of our study. When testing sensitivity to COM parameter values, we performed this analysis as an elasticity analysis, changing individual parameter values by a set percentage and examining the resulting percent changes in the results. When testing sensitivity to the total removals reconstruction, we examined results of alternative reconstructions.

**COM parameter elasticity analysis**

We performed the elasticity analysis by changing the value of a single model parameter by 50%, and then running all COMs and the superensemble models with this changed value, leaving all other inputs unchanged. We then looked at the resulting percent change of each COM and superensemble model prediction of the mean and slope of $B/{B_{\mathrm{MSY}}}$ of the last 5 years of data. If the percent change of a model result exceeded 50% (the percent change in the parameter value), the model was considered to be sensitive to changes in that parameter.

Mean $B/{B_{\mathrm{MSY}}}$ results appeared to not be sensitive to changes in COM parameters (Figure D1), as no model result of mean $B/{B_{\mathrm{MSY}}}$ exceeded 50% change from the base run. The greatest percent change was observed for the final biomass range.

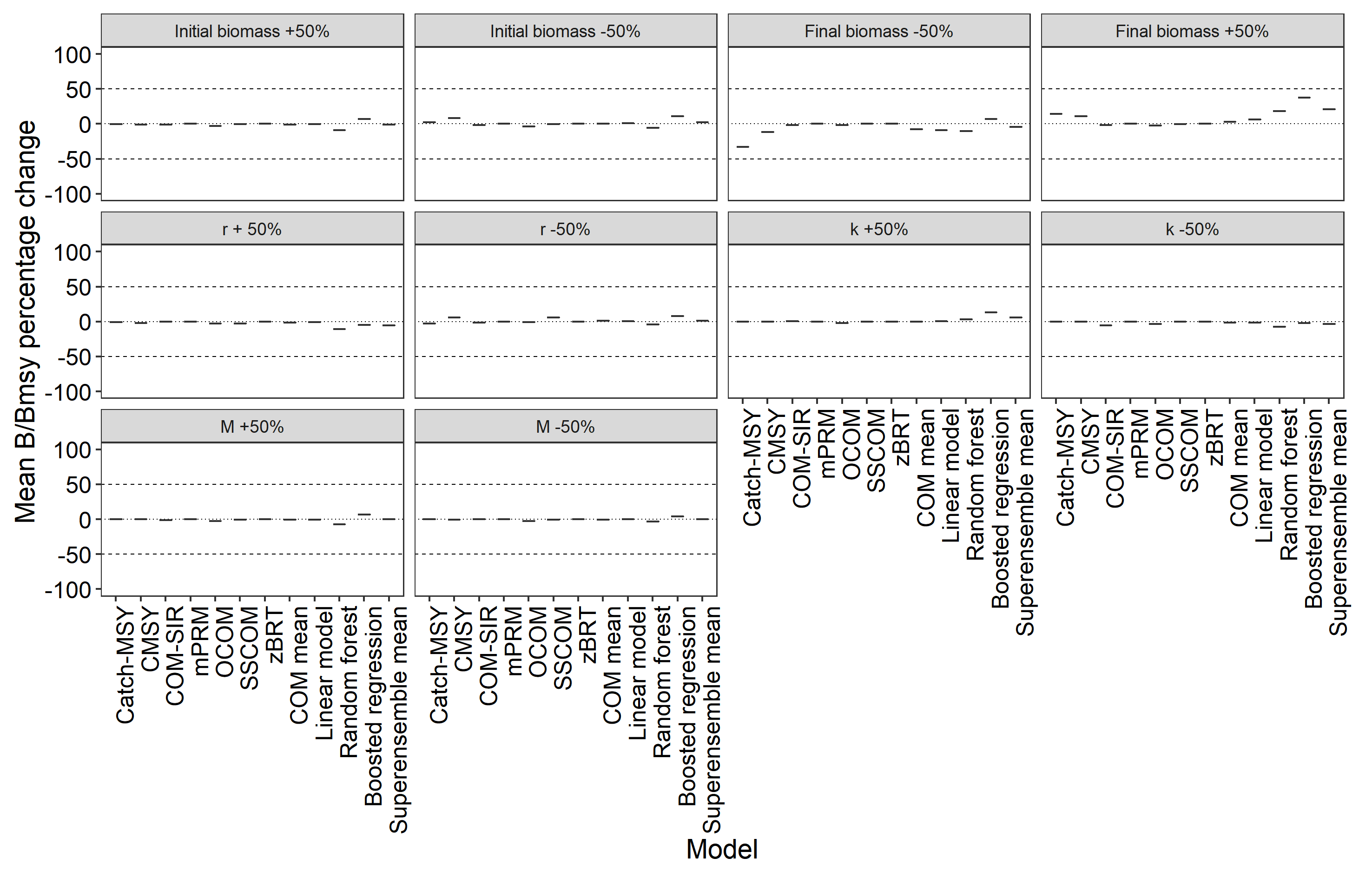

Figure D1: Elasticity of model results (mean $\boldsymbol{B}/{\boldsymbol{B}_{\mathbf{MSY}}}$ over the last 5 years of data) to changes in COM parameters. The horizontal dashed lines indicate changes of -50% and 50% when compared to the base run as described in the main text.

$B/{B_{\mathrm{MSY}}}$ slope results were sensitive to changes in multiple COM parameters (Figure D2). Catch-MSY results were sensitive to a higher final biomass range, and COM-SIR results were sensitive to both positive and negative changes in the initial biomass range, *r*, as well as negative changes in *k* and *M*. However, none of the superensemble estimates of $B/{B_{\mathrm{MSY}}}$ slope were sensitive to changes in COM parameters, with the greatest percent changes being observed for increases in the final biomass range and *r*.

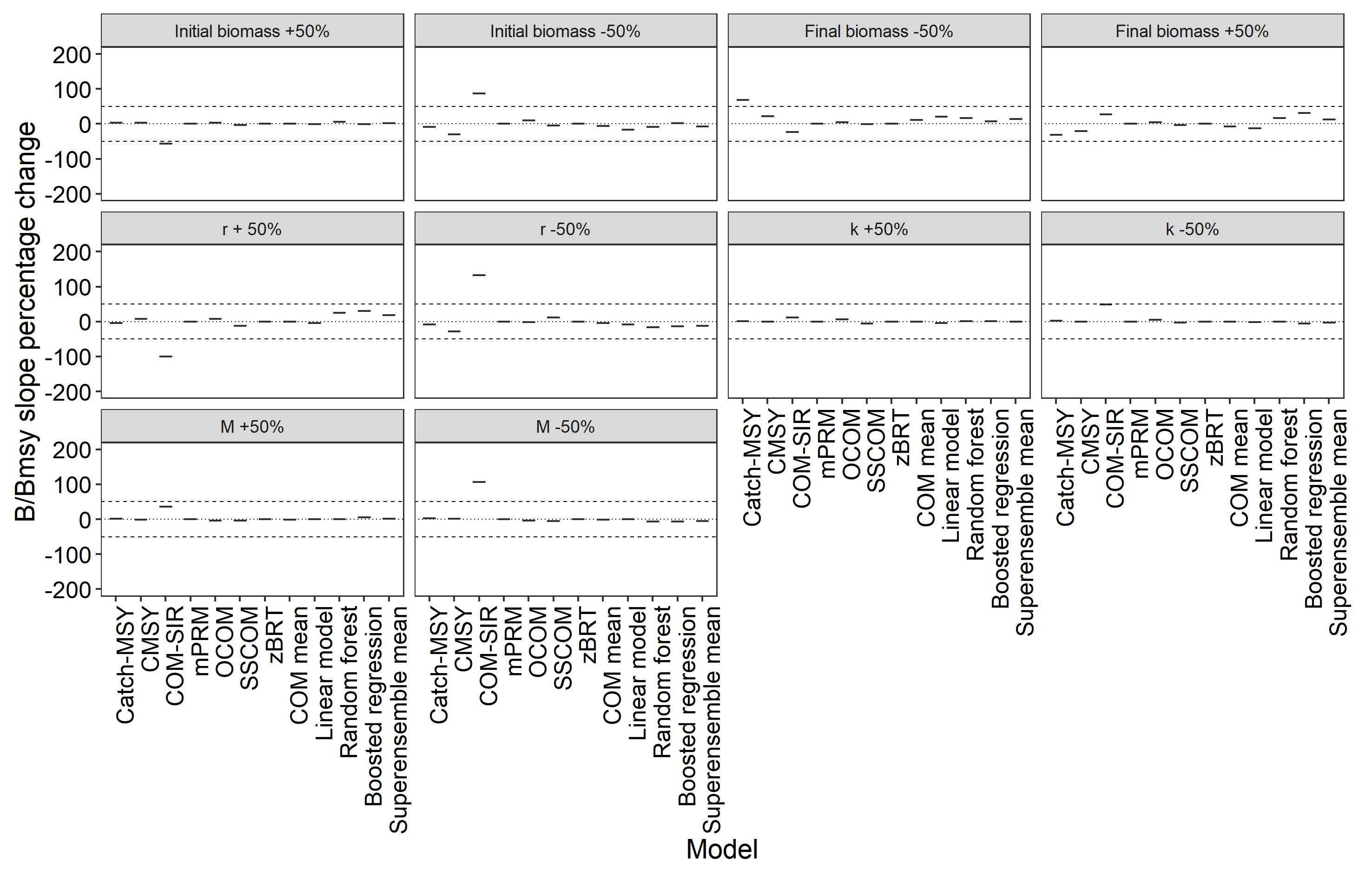

Figure D2: Elasticity of model results ($\boldsymbol{B}/{\boldsymbol{B}_{\mathbf{MSY}}}$ slope over the last 5 years of data) to changes in COM parameters. The horizontal dashed lines indicate changes of -50% and 50% when compared to the base run as described in the main text.

**Total removals time-series sensitivity analysis**

Several assumptions were made when reconstructing the time-series of total pike removals. To test for the models’ sensitivity to these assumptions, we reconstructed the total pike removals time-series under various alternative assumptions, and then ran the COMs and superensemble models with each new time-series, examining the percent change of each COM and superensemble model prediction of the mean and slope of $B/{B_{\mathrm{MSY}}}$ of the last 5 years of data. For this, we made a total of 17 alternative reconstructions of total pike removals. These reconstructions are detailed below.

*Reconstruction 1*

The models’ sensitivity to consistent overestimation of the commercial removals time-series was tested. For this, annual removals from the commercial removals time-series were reduced with 50%.

*Reconstruction 2*

The models’ sensitivity to consistent underestimation of the commercial removals time-series was tested. For this, annual removals from the commercial removals time-series were increased with 50%.

*Reconstruction 3*

The models’ sensitivity to consistent overestimation of the recreational removals time-series was tested. For this, annual removals from the recreational removals time-series were reduced with 50%.

*Reconstruction 4*

The models’ sensitivity to consistent underestimation of the recreational removals time-series was tested. For this, annual removals from the recreational removals time-series were increased with 50%.

*Reconstruction 5*

The models’ sensitivity to the assumption that the number of commercial fishing vessels registered in the Rügen region remained constant before 1991 was tested. For this, we reconstructed the recreational removals time-series under two alternative assumptions. The first assumption was that commercial fishing vessels experienced a linear decline in the years leading up to 1991, calculated by fitting a simple linear regression model to the 1992-2018 data on registered commercial fishing vessels in the Rügen region, and extrapolating back in time to 1955 (Figure D3a).

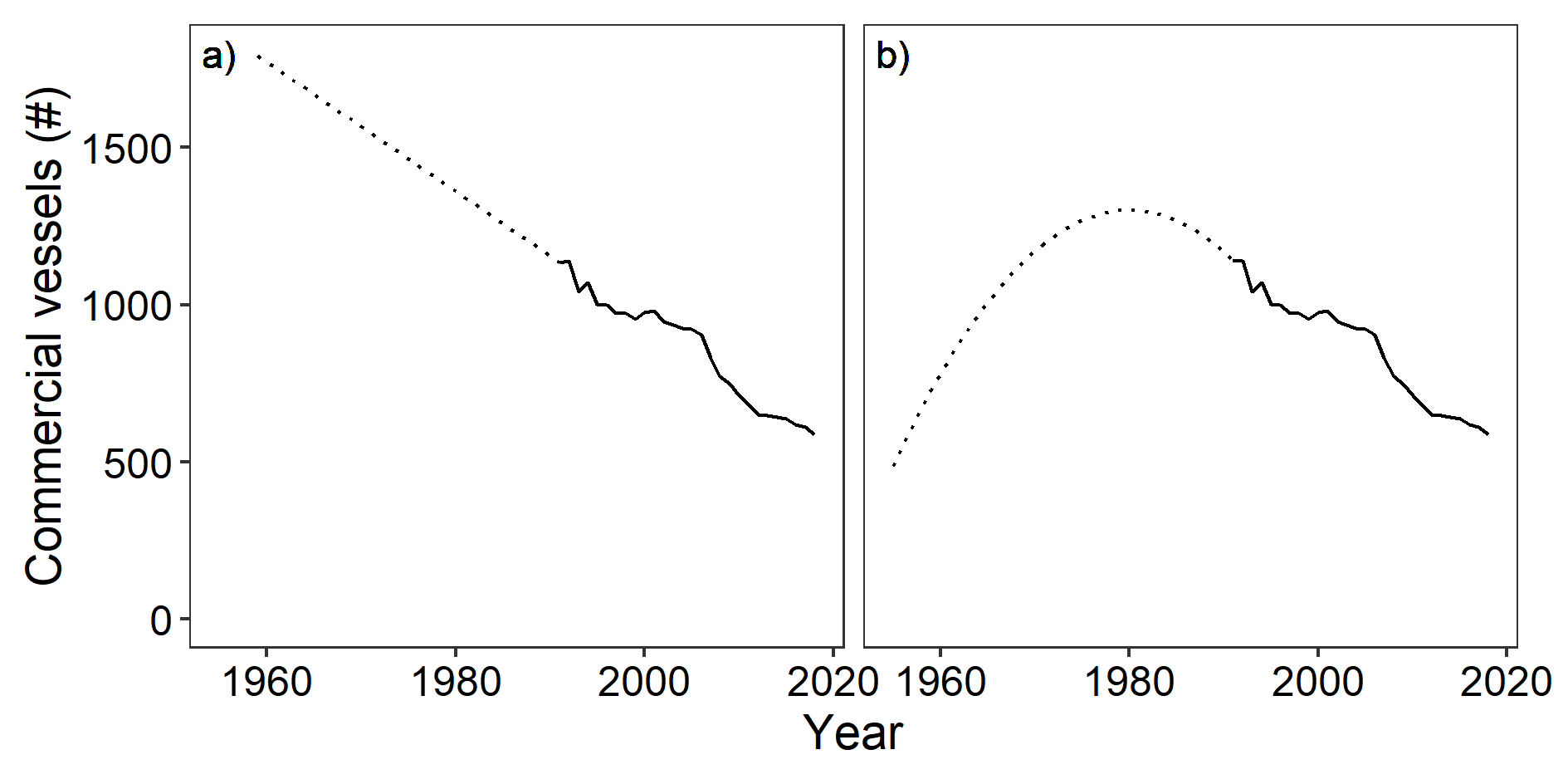

Figure D3: Alternative reconstructions of the number of commercial fishing vessels registered in the Rügen area before 1992, assuming a linear decline in vessel numbers (a), or an initial increase and subsequent decrease in vessel numbers (b).

*Reconstruction 6*

Following Reconstruction 5, the second assumption was that the number of commercial fishing vessels increased from 1955 to 1980, and subsequently started declined, following a parabolic shape (Figure B1b).

*Reconstruction 7*

Same as the main text reconstruction, except that the resident harvest and release rate is calculated as the average of the 2007 and 2015 data.

*Reconstruction 8*

Same as Reconstruction 7, except that the resident CPUE constant of proportionality is determined from the 2015 angling catches instead of the average of 2007 and 2015, and that the resident harvest and release rate is determined only from the 2015 data.

*Reconstruction 9*

Same as reconstruction 8, except that the CPUE constant of proportionality is determined from the 2015 total angling removals instead of the catches. Based on this constant, a single time-series of recreational removals is recreated for the entire time-period 1955-2018, instead of first reconstructing separate time-series for resident and tourist recreational removals for 1992-2018, and then extrapolating angler removals back to 1955.

*Reconstruction 10*

Instead of linearly extrapolating resident trips per annual coastal angling license between 2007 and 2015, resident trips per annual coastal angling license were calculated as the mean of the 2007 and 2015 estimate. Using this mean and the number of annual coastal angling licenses for the other years, a time-series of resident angling trips to the Rügen area was extrapolated for the years 1991-2018. 2015 tourist angling trips were calculated using the new estimate for 2015 resident angling trips, and the observed ratio of 2015 resident to tourist angling trips from (Weltersbach et al., in press). Using this new estimate of 2015 tourist angling trips, together with the time-series of short-term coastal angling licenses, a time-series of tourist angling trips was extrapolated for 1991-2018. Adding the resident angling trips gave a time-series of total angling trips to the Rügen area for 1991-2018. Then, the number of angling trips in 1990 was estimated by linearly extrapolating the total angling trips from 1991-2000 back to 1990. Next, using the DAV membership data from 1954-1990 the 1990 ratio between Rügen angling trips and DAV membership was calculated. Then, by assuming this ratio remained constant, the number of angling trips was extrapolated back to 1955. Next, using the 2015 recreational harvest and release rate (Weltersbach et al., in press) and 2015 number of angling trips, total recreational removals were calculated for 2015, as well as the 2015 recreational CPUE. Using 2015 recreational and commercial CPUE, a CPUE constant of proportionality was calculated. This CPUE constant of proportionality was used together with the time-series of commercial CPUE and recreational effort (number of trips) to extrapolate recreational removals for 1955-2018.

*Reconstruction 11*

Same as Reconstruction 10, but the CPUE constant of proportionality was calculated from the total 2015 recreational catch, instead of recreational removals. For this, a separate CPUE constant of proportionality was calculated for recreational and tourist catches. Individual 1991-2018 time-series of recreational and tourist removals were thus calculated first, then added together. Then, for the reconstruction of pre-1990 recreational removals, recreational CPUE was first calculated for 1991. From this, a CPUE constant of proportionality was calculated for 1991. This constant, together with the 1955-1990 reconstructed time-series of Rügen angling trips and the 1955-1990 commercial CPUE time-series, was used to extrapolate total angler removals back to 1955.

*Reconstruction 12*

Same as reconstruction 11, but the resident CPUE constant of proportionality was calculated as the average of the 2007 and 2015 CPUE constant of proportionality. Furthermore, the resident harvest and release rate was calculated as the average between 2007 and 2015.

*Reconstruction 13*

Same as reconstruction 12, but the resident harvest and release rate between 2007 and 2015 was interpolated as a linear relationship between the two years, extrapolated backward to 1991 as constant to 2007, and extrapolated forward to 2018 as constant to 2015.

*Reconstruction 14*

The recreational removals time-series was reconstructed by using only the data from the 2015 telephone-diary study (Weltersbach et al., in press), and ignoring the data from the 2007 telephone-diary study (Dorow & Arlinghaus, 2011). Using the 2015 estimate of Rügen resident angling trips and the 1991-2018 time-series of annual coastal angling licenses, a 1991-2018 time-series of resident angling trips was extrapolated by assuming a constant proportion of resident angling trips per annual coastal angling license. Similarly, a 1991-2018 time-series of tourist angling trips was reconstructed by using the 2015 estimate of Rügen tourist angling trips and the 1991-2018 time-series of short-term coastal angling licenses. Then, the number of angling trips in 1990 was estimated by linearly extrapolating the total angling trips from 1991-2000 back to 1990. Next, using the DAV membership data from 1954-1990, the 1990 ratio between Rügen angling trips and DAV membership was calculated. Then, by assuming this ratio remained constant, the number of angling trips was extrapolated back to 1955. Next, recreational CPUE was estimated for 2015 from the 2015 estimate of total angler removals and the 2015 estimate of total number of angling trips. Using the 2015 recreational and commercial CPUE, a CPUE constant of proportionality was calculated. Using this constant and the reconstructed 1955-2018 time-series of recreational effort, recreational removals were reconstructed for 1955-2018.

*Reconstruction 15*

Same as Reconstruction 14, but the CPUE constant of proportionality was calculated from the total 2015 recreational catch, instead of recreational removals. For this, a separate CPUE constant of proportionality was calculated for recreational and tourist catches. Individual 1991-2018 time-series of recreational and tourist removals were thus calculated first, then added together. Then, for the reconstruction of pre-1990 recreational removals, recreational CPUE was first calculated for 1991. From this, a CPUE constant of proportionality was calculated for 1991. This constant, together with the 1955-1990 reconstructed time-series of Rügen angling trips and the 1955-1990 commercial CPUE time-series, was used to extrapolate total angler removals back to 1955.

*Reconstruction 16*

Same as reconstruction 15, but the resident CPUE constant of proportionality was calculated as the average of the 2007 and 2015 CPUE constant of proportionality. Furthermore, the resident harvest and release rate was calculated as the average between 2007 and 2015.

*Reconstruction 17*

Same as reconstruction 16, but the resident harvest and release rate between 2007 and 2015 was interpolated as a linear relationship between the two years, extrapolated backward to 1991 as constant to 2007, and extrapolated forward to 2018 as constant to 2015.

**Results**

Mean $B/{B_{\mathrm{MSY}}}$ results of the models differed from their base run results to a varying degree for the different reconstructions of total removals (Figure D4), although most remained largely unaffected for most reconstructions. Notably, 50% changes in the time-series values of either commercial or recreational removals (Reconstructions 1-4), as well as different assumptions regarding pre-1991 commercial effort (Reconstructions 5-6), did not have a large impact on the final superensemble estimate of mean $B/{B_{\mathrm{MSY}}}$. Reconstruction 13 seemed to result in the largest percent change of mean $B/{B_{\mathrm{MSY}}}$ of the superensemble models.

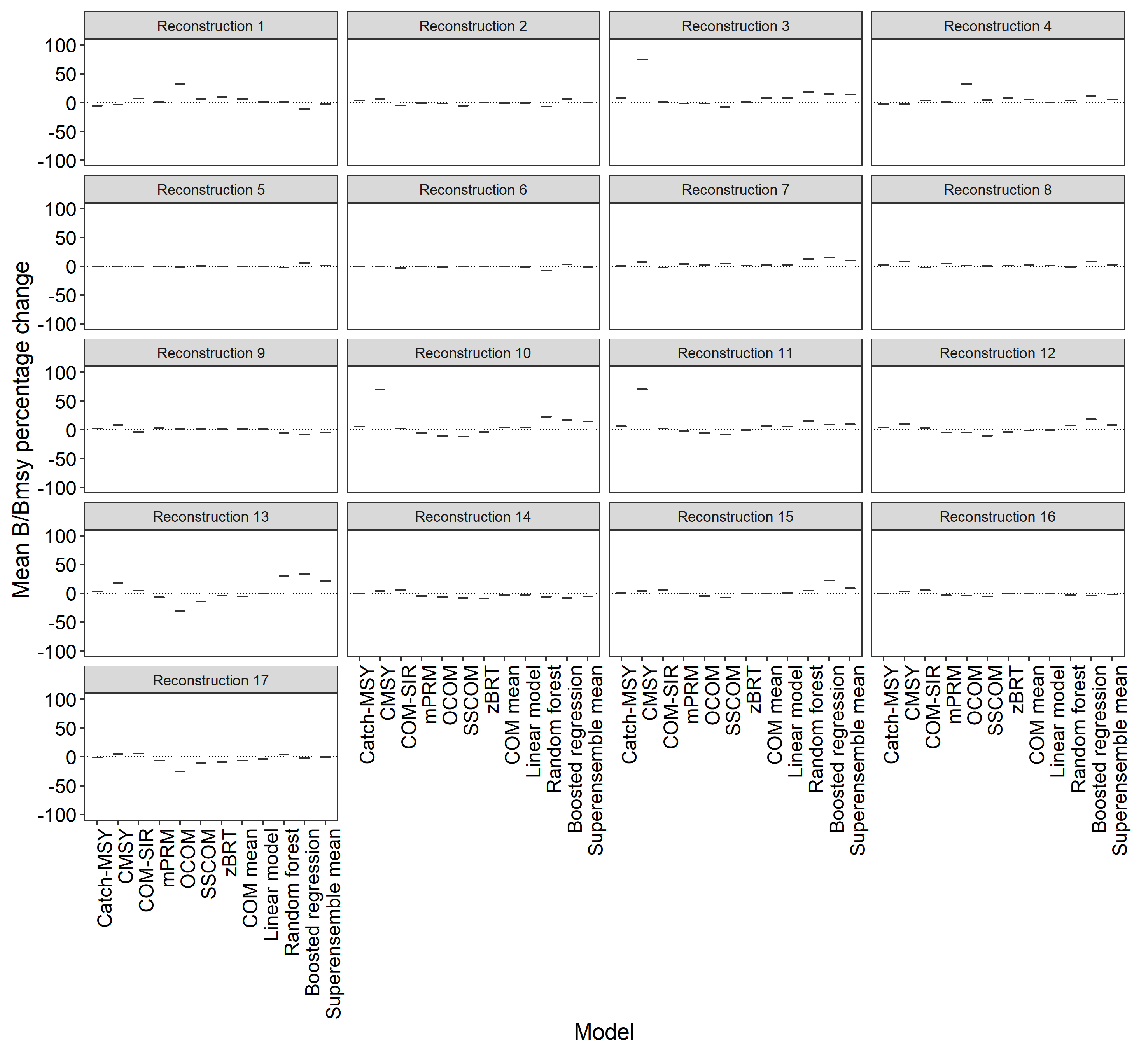

Figure D4: Sensitivity of model results (mean $\boldsymbol{B}/{\boldsymbol{B}_{\mathbf{MSY}}}$ over the last 5 years of data) to different reconstructions of total removals of pike, shown as percent change when compared to the base run as described in the main text.

$B/{B_{\mathrm{MSY}}}$ slope results of the models differed from their base run results to a varying degree for the different reconstructions of total removals (Figure D5). Notably, 50% changes in the time-series values of either commercial or recreational removals (Reconstructions 1-4), as well as different assumptions regarding pre-1991 commercial effort (Reconstructions 5-6), did not have a large impact on the final superensemble estimate of the $B/{B_{\mathrm{MSY}}}$ slope. COM-SIR appeared to be the most sensitive to the different reconstructions, possibly resulting from its base run slope being close to 0.

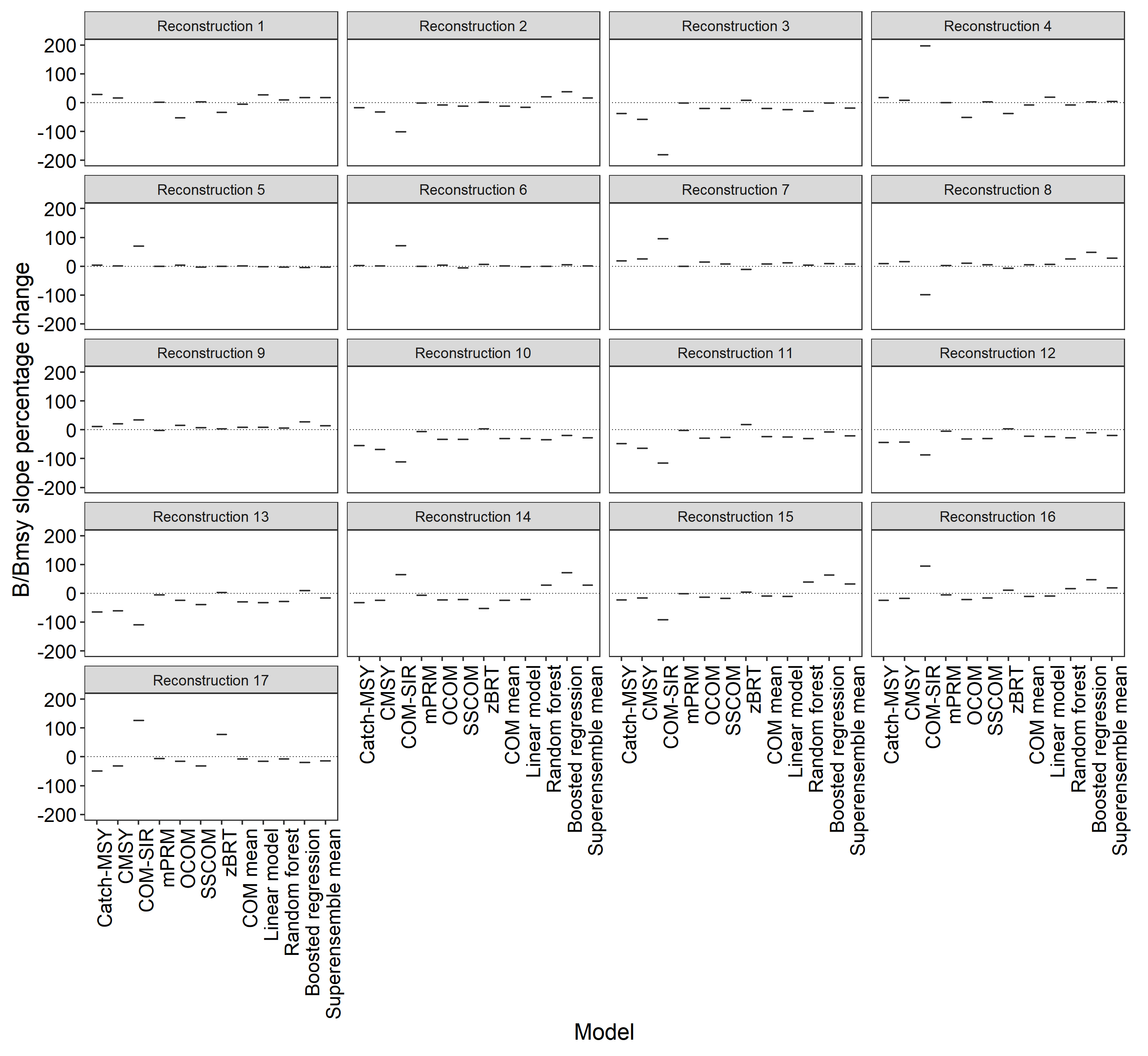

Figure D5: Sensitivity of model results ($\boldsymbol{B}/{\boldsymbol{B}_{\mathbf{MSY}}}$ slope over the last 5 years of data) to different reconstructions of total removals of pike, shown as percent change when compared to the base run as described in the main text.

Weltersbach, M. S., Riepe, C., Lewin, W. C., & Strehlow, H. V. (in press). *Ökologische, soziale und ökonomische Dimensionen des Meeresangelns in Deutschland*. Thünen-Institut für Ostseefischerei.
